# Using wingbeat frequency to estimate mass gained by seabirds

**DOI:** 10.1101/2025.05.15.654001

**Authors:** Allison Patterson, Marie Auger-Méthé, Kyle Elliott

## Abstract

1. Energy intake is a fundamental currency in ecology that is critical to reproductive success, survival and lifetime fitness. Measuring foraging success in wild animals via biologgers has been a long-standing challenge.

2. Flying animals gain mass during foraging, and they must counteract the associated increased gravitational force by creating additional lift. Pennycuick (1996) proposed that wingbeat frequency (*w*) should vary with the square root of body mass 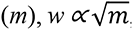, when other variables influencing wingbeat frequency are held constant.

3. We present a state-space model that estimates instantaneous changes in body mass by modelling this relationship with wingbeat frequency using animal-borne accelerometer and depth data. To demonstrate the usefulness of our proposed method, we applied it to biologging data from 55 thick-billed murres (*Uria lomvia*) during the incubation period.

4. Our mass estimates allowed us to identify areas associated with higher gains, and to demonstrate that foraging success was generally higher farther from the colony. However, 79% of foraging trips were associated with mass deficits. We performed simulation studies to assess the sensitivity of our method to parameter misspecification and the increase in accuracy gained from including the known mass at recapture. As estimates of energy intake allow for testing of long-standing hypotheses in foraging ecology, our method provides a new tool to help answer these questions with any animal that engages in flapping flight.

## Introduction

Understanding how animals manage their energy and time budgets is a fundamental goal of foraging theory (Charnov, 1976; MacArthur & Pianka, 1966; Orians & Pearson, 1979; Stephens & Krebs, 1986). Balancing energy expenditure and energy intake is critical to reproductive success, survival, and lifetime fitness. To test many long-standing hypotheses about foraging behaviour, ecologists require fine-scale continuous measures of energy intake in free-living animals (Schick et al., 2013). However, methods for measuring energy intake in wild animals have remained a long-standing challenge for ecologists.

Biologgers are now able to accurately measure energy expenditure, via accelerometry (Elliott, Le Vaillant, et al., 2013; Wilson et al., 2020) or heart-rate logging (Butler et al., 2004; Green, 2011; Hicks et al., 2017). Energy intake has been more challenging to estimate via biologgers, but several recent studies have used accelerometers to measure prey capture rates based on rapid changes in acceleration, direction or posture that likely indicate prey capture attempts (Chimienti et al., 2016; Clermont et al., 2021). Ideally, algorithms for detecting prey capture from animal movements would be validated with independent measurements such as beak openings (Brisson-Curadeau et al., 2021), stomach temperature loggers (Ropert-Coudert & Kato, 2006; Wilson et al., 1995), or video recordings (Lok et al., 2023; Sidrow et al., 2024; Watanabe & Takahashi, 2013); however, these types of validation data are difficult to obtain in the wild, especially for smaller species. Many methods exist to identify foraging areas based on movement tracks (i.e., time series of geographic locations), but relying solely on spatial movement patterns to infer prey capture can lead to erroneous assumptions about important foraging areas (Bennison et al., 2018; Florko et al., 2023).

Biomechanical approaches estimating buoyancy from dive drift rates have been used to estimate changes in body condition of marine mammals and sharks using time-depth recorders (Biuw et al., 2003, 2007; Del Raye et al., 2013; Robinson et al., 2010; Yong et al., 2024). This approach assumes that increases in an animal’s lipid stores will result in increased buoyancy while swimming, which in turn can be measured based on the drift rate when an animal is floating passively in the water column (Biuw et al., 2003). Biomechanical approaches have been used to estimate mass gain patterns in juvenile elephant seals (Biuw et al., 2003) and map spatial variation in foraging profitability of elephant seals in the Southern Ocean (Biuw et al., 2007). Migrating white sharks showed consistent increases in drift rate, indicating depletion of lipid stores to fuel migration (Del Raye et al., 2013). This approach was further developed into a state-space model that estimates a continuous time series of body condition where lipid gain can be modelled as a function of environmental and behavioural characteristics (Schick et al., 2013). Unfortunately, the drift rate technique is only applicable to some diving species. The buoyancy of diving seabirds is also influenced by variation in the air volume in their feathers and lungs, which likely overwhelms any variation in buoyancy from lipid density (Kooyman & Ponganis, 1998; Lovvorn et al., 2004). Moreover, as it takes days to convert food into lipids, this technique can only provide information over longer time scales and for larger species that store significant energy as lipids.

For flying animals, Pennycuick (1996) proposed that wingbeat frequency (*w*) should vary with the square root of body mass 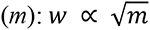. The effects of feeding and consuming fuel in daily activities result in changes in mass that will alter wingbeat frequency while most other factors influencing wingbeat frequency (e.g. wing span, wing area, and wing moment of inertia) would remain constant (Pennycuick, 1996). As an individual increases their mass during foraging, the body needs to counteract that increased gravitational force by creating additional lift. At a fixed forward speed and body shape, this can only be done by altering the angle of attack or increasing the wingbeat frequency or amplitude. At high speeds in wind tunnels, it is primarily the wingbeat frequency that is varied in response to body mass (Hambly et al., 2004).

The relationship between mass and wingbeat frequency was first demonstrated in free-living birds with European shags, *Phalacrocorax aristotelis* (Sato et al., 2008). Using accelerometers, Sato et al (2008) estimated changes in mass based on the change in mean wing beat frequency between bouts of flying. They found a strong relationship between estimated change in mass and both dive bout duration and cumulative dive duration within bouts. Estimated mass loss while on land (based on the difference between the previous in-bound flight and subsequent out-bound flight) was positively related to mass gain from the previous trip but not to total time spent on land. The advantage of this biomechanical approach over other accelerometer-based approaches is that it gives an estimate of mass gained over time rather than prey capture attempts.

We developed a state-space model that estimates changes in mass for diving seabirds based on measurements of wingbeat frequency and foraging behaviour measured with accelerometer and time-depth recorders. This represents a significant advancement in the approach of Sato et al (2008), which allows for continuous estimates of mass; the potential to include covariates that influence mass change; and the ability to account for measurement error in wing beat frequency observations. The process equation of our model estimates continuous changes in mass based on mass loss through regular metabolic activity, increased energy expenditure in flight, and mass gained while diving. The observation equation is based on Pennycuick’s equation for the relationship between wingbeat frequency and mass. The state-space model can estimate the effect of covariates on mass gain while foraging. We illustrate the usefulness of our model with a case study using tracking data collected from thick-billed murres (hereafter murres, *Uria lomvia)*. Murres are a pursuit diving seabird species that uses constant flapping-flight and has exceptionally high wing-loading (Elliott, Le Vaillant, et al., 2013; Elliott, Ricklefs, et al., 2013). We also perform simulation studies to assess the changes in accuracy of our model estimates under various conditions.

## Methods

### State-Space Model

Our state-space model is a hierarchical model with three levels. The first level, often referred to as the process equation (Auger-Méthé et al., 2021), models the mass of individual *i* at time *t*, 𝑚_𝑖,𝑡_, as a function of its previous mass and behaviour (resting, flying, or diving):

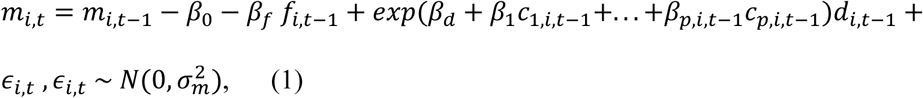

where the mass of individual *i* at time *t,* 𝑚_𝑖,𝑡_, can be considered the hidden state of the state-space model. The model contains two indicator variables, one which specifies whether the individual is flying at time *t* (𝑓_𝑖,𝑡_ = 1) and the other specifies whether it is diving (𝑑_𝑖,𝑡_ = 1). The baseline rate of mass loss an individual incurred during the time interval 𝛥𝑡 is represented by 𝛽_0_, while 𝛽_𝑓_ represents the additional rate of mass loss incurred while flying. To ensure that these capture mass loss, the coefficients are forced to be greater than 0 (𝛽_0_ > 0, 𝛽_𝑓_ > 0). As diving in murres generally does not incur much higher energy expenditure than the baseline metabolic rate (Elliott, Ricklefs, et al., 2013), we assumed that diving can only be associated with a relative increase in mass compared to the baseline rate of mass loss. We model this relative mass gain using an exponential function of *p* covariates, 𝑐_1_, . . ., 𝑐_𝑝_, which is similar to using a log-link function in a generalized linear model (Zuur, 2007). This function has an intercept-like parameter, 𝛽_𝑑_, which represents the increase in mass when all covariates are equal to 0, with negative values with a large magnitude (e.g., -100) being interpreted as negligible relative gain in mass. The remaining coefficients, 𝛽_1_, . . ., 𝛽_𝑝_, represent how the covariates modify the rate of gain while diving. These coefficients are not forced to be greater than 0, and negative values will represent decreases in relative gain in mass with increased covariate values. The process stochasticity, 𝜖_𝑖,𝑡_, allows for unexplained variation in this relationship and is modelled with a Normal distribution centered at 0 and with variance 𝜎^2^ .

The second level of the state-space model, often referred to as the observation equation (Auger-Méthé et al., 2021), represents the relationship between the mass of the individual and its wingbeat frequency at time *t*:

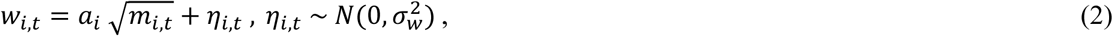

where the observation, 𝑤_𝑖,𝑡_, represents an individual’s wingbeat frequency at time *t* and 𝑎_𝑖_ represents an individual-specific constant for the relationship between *w* and *m* for individual *i*. The measurement error at time *t*, 𝜂_𝑖,𝑡_, is modelled with a Normal distribution centered at 0 and with variance 𝜎^2^. The constant 𝑎_𝑖_ accounts for differences in physical aspects such as wing span and wing area, sometimes modelled independently (Pennycuick, 1996). As such, the third level of the state-space model allows for the relationship between the mass and the wingbeat frequency to change between individuals:

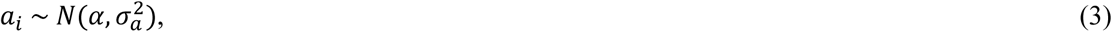

which can be thought of as a random effect modelled with a Normal distribution centered at the mean population value of this constant, 𝛼, and with variance 𝜎^2^.

This state-space model could be applied to data using various inferential frameworks and computational tools (for examples, see Auger-Méthé et al. 2021), but as exemplified with the case study and simulation study, we used a maximum likelihood framework and the R package TMB (Kristensen et al., 2016). Under this framework, all coefficients from equation 1 (𝛽_0_, 𝛽_𝑓_, 𝛽_𝑑_, 𝛽_1_, . . ., 𝛽_𝑝_), 𝛼 of equation 3, as well as up to the three standard deviations (𝜎_𝑚_, 𝜎_𝑤_, 𝜎_𝑎_) must be estimated (see *Simulation studies* section for a discussion of potential estimation issues that may arise if all three are estimated simultaneously). The mass of individual *i* at time *t*, 𝑚_𝑖,𝑡_, and the individual-specific values of 𝑎_𝑖_ are predicted. We used R version 4.3.1 (R Core Team, 2023) and TMB version 1.9.15 (Kristensen et al., 2016).

### Case study

Tracking data used in the case study were collected from 55 biologger deployments on murres breeding at Coats Island, NU (82.01° W, 52.95° N) in 2023. In total, deployments were conducted on 52 unique murres and three murres had two deployments each. Murres incubating eggs were captured at the nest using a noose pole. We attached AxyTrek Mini (Technosmart, Rome, Italy) loggers to the feathers on the lower back using Tesa Tape, cable ties, and super glue. Loggers were programmed to collect GPS (1 min), depth (1 Hz), and tri-axial acceleration (50 Hz). Murres were recaptured 12-48 hours later for device retrieval. Each murre was weighed to the closest gram at initial deployment and recapture. All animal handling was conducted under permit.

Accelerometers can be used to measure the dominant stroke frequencies of animals while flying and swimming (Sato et al., 2007). Wingbeat frequencies (*w*) were calculated from the heave (z) axis of accelerometers by identifying the frequency with the highest power from a fast Fourier transform calculated over a 10-second moving window (Patterson et al., 2019). To focus on signals associated with flight, time steps with an interquartile range <0.1 gravity were assigned a *w* of 0 Hz. Pitch was calculated from the static component of the surge (x), sway (y), and heave (y) axes smoothed over a 1- sec moving window (Shepard, Wilson, Quintana, et al., 2008). Before calculating pitch, each axis was centred at 0 gravity during periods of presumed flight (6.5 Hz > *w* < 9.0 Hz), this was to account for differences in tag placement among birds, so that pitch during flight was standardized to 0 across deployments (Shepard, Wilson, Halsey, et al., 2008).

We used the biologger data to classify tracks into four basic behaviours at 10-sec intervals: flying, diving, resting at colony, and resting on water. We used a two-component mixture model with a Weibull distribution to preliminarily assign observations to flying based on *w* (flying = 7.83 ± 0.54 [mean ± sd], not flying = 0.52 ± 0.80). Observations were retained as flying if they were part of 6 consecutive observations (e.g. 60-sec) of flying. Observations with maximum depth >1 m were assigned a behaviour of diving. We used a continuous-time correlated random walk model, implemented with the aniMotum package (Jonsen et al., 2023), to interpolate GPS locations at 10-sec. For each interpolated location, we calculated the distance between the location estimate and the colony. Unclassified observations within 500 m of the colony were classified as resting at colony and remaining observations were classified as resting on water.

We included two covariates for the gain rate while diving in the state-space model: maximum diving depth (c_depth_) and standard deviation of pitch (c_pitch_). Maximum dive depth was calculated separately for each dive and this value was assigned to all 10-sec intervals within that dive. Standard deviation of pitch was calculated over each 10-sec interval. The ranges of both covariates were scaled to values between 0-1 by dividing by the maximum observed value. The case study model included a random effect of individual on *a*, the coefficient that scales the relationship between wingbeat frequency and mass. We fitted the model described in equations 1-3 using the R package TMB, and to resolve convergence issues we fixed the value of 𝜎_𝑚_ to 0.1 (see *Simulation studies* section for a sensitivity analysis for this parameter). We included the observed initial mass and final mass for each deployment as fixed values in the model, forcing mass estimates to conform to the known mass at these times. We also refit the model without final mass and calculated the root mean squared error (RMSE) between observed and estimated final mass as a measure of how much estimates of final mass deviated from true values.

We defined foraging trips as any movement away from the colony that lasted longer than 4 hours and included at least 15 minutes of diving. We excluded one unusually long trip that lasted 32.7 hr, which was more than twice as long as the next longest trip (15.6 hr). We recorded 48 foraging trips from 46 deployments, in nine deployments the murres remained at the colony the entire time and two murres recorded multiple trips during a deployment. For each trip, we calculated the trip duration (hr), maximum distance from colony (km), total distance (km), time spent in each behaviour (hr), net mass gain (g), and gross mass gain (g). We used linear regression to examine the influence of trip duration, maximum distance from the colony, and starting mass on net gain during a foraging trip.

We used the estimated gain rate across all tracks to visualize the energetic landscape around the colony. First, we summarized the average gain rate while diving at 3-hr intervals. Only intervals with at least 15 min of diving were included in this analysis. For each 3-hr diving interval, we calculated the median latitude and longitude while diving. We then applied a spatial smoothing function using the ‘spatstat’ R package, version 3.1-1 (Baddeley et al., 2016). The analysis used a 5 km smoothing bandwidth on a 1 x 1 km grid. Raster cells with no foraging observations within 10 km were masked from the analysis. We report the smoothed estimates with standard errors to show regions with higher uncertainty. All code used in the case study is available in Supplemental Materials.

### Simulation studies

We performed two simulation studies. The first explored how misspecifying the value of the standard deviation of the process equation (𝜎_𝑚_) affects the model accuracy, while the second explored how including the mass of the individual at recapture improves the model. State-space models can have difficulty simultaneously estimating the standard deviations associated with process stochasticity and measurement error (Auger-Méthé et al., 2016, 2021), and when we tried to do so with the murre data, we had convergence problems. Thus, we decided to fix the value of 𝜎_𝑚_ to 0.1. While we chose a biologically relevant value, the exact value chosen is arbitrary. Our first simulation study is a sensitivity analysis that assesses the robustness of our results to the misspecification of 𝜎_𝑚_.

The second simulation study explored whether having known values for some of the hidden states, sometimes referred to as a semi-supervised approach (Saldanha et al., 2023; Sidrow et al., 2024), can improve estimation of the state-space models. As is relatively common in tagging studies of birds where the tagged individual is recaptured to recover the biologger, we weighed each murre at capture and recapture. Thus, we included both initial and final mass in the model. As some studies do not recapture individuals, because data are acquired through remote downloads, we wanted to explore the effect of including the final mass in the model. Both simulation studies also assess whether the model has estimatability issues, which is an important aspect of model checking when using state-space models (Auger-Méthé et al., 2016, 2021).

The base simulation structure for both studies was the model defined in equations 1-3 using the estimated parameter values from the case study (Table 1). To create simulations similar to real data, we use the time series of observed covariates (max depth of a dive: 𝑐_𝑑𝑒𝑝𝑡ℎ,𝑖,𝑡_; standard deviation of pitch: 𝑐_𝑝𝑖𝑡𝑐ℎ,𝑖,𝑡_) and of the indicator variables (flying: 𝑓_𝑖,𝑡_ ; diving: 𝑑_𝑖,𝑡_) from the murre dataset, as well as the observed mass of the birds at the start and end of the time series. For each simulation, we randomly selected 20 tracks from the 55 murre tracks used in the case study and we ran 100 simulations per scenario. Because we used the covariate and indicator time series from the murre dataset, the length of the time series varied. For the first simulation study, we changed the value of 𝜎_𝑚_ used to create the simulations and to fit the model (i.e., fixed 𝜎_𝑚_). The 𝜎_𝑚_ values explored were: (0.05, 0.1, 0.15). As we looked at all combinations of simulated and fixed 𝜎_𝑚_ values, we explored a total of nine different scenarios in the first simulation study. For the second simulation study, we used 𝜎_𝑚_ = 0.1, as in the case study, and for each of the 100 simulations, we fitted two models: one which included the final mass (i.e., mass at recapture), and another that ignored the final mass value.

**Table 1.**
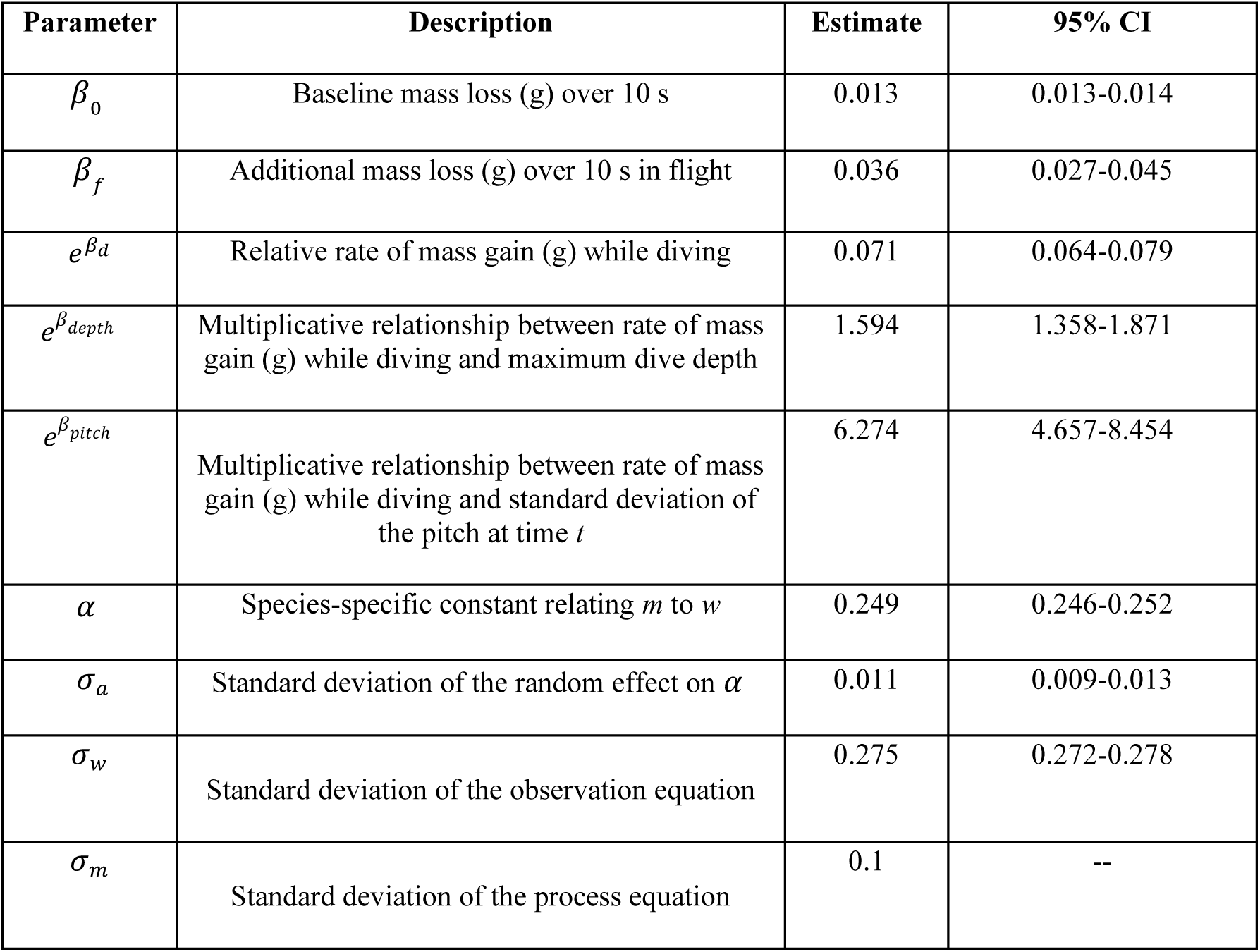
Parameter estimates from the state-space model estimating changes in mass based on wingbeat frequency for incubating thick-billed murres from Coats Island, NU. Note that the parameters associated with diving are exponentiated for ease of interpretation (see equation 1).

| Parameter | Description | Estimate | 95% CI |
| --- | --- | --- | --- |
| $\beta_0$ | Baseline mass loss (g) over 10 s | 0.013 | 0.013-0.014 |
| $\beta_f$ | Additional mass loss (g) over 10 s in flight | 0.036 | 0.027-0.045 |
| $e^{\beta_d}$ | Relative rate of mass gain (g) while diving | 0.071 | 0.064-0.079 |
| $e^{\beta_{depth}}$ | Multiplicative relationship between rate of mass gain (g) while diving and maximum dive depth | 1.594 | 1.358-1.871 |
| $e^{\beta_{pitch}}$ | Multiplicative relationship between rate of mass gain (g) while diving and standard deviation of the pitch at time $t$ | 6.274 | 4.657-8.454 |
| $\alpha$ | Species-specific constant relating $m$ to $w$ | 0.249 | 0.246-0.252 |
| $\sigma_a$ | Standard deviation of the random effect on $\alpha$ | 0.011 | 0.009-0.013 |
| $\sigma_w$ | Standard deviation of the observation equation | 0.275 | 0.272-0.278 |
| $\sigma_m$ | Standard deviation of the process equation | 0.1 | -- |

For both simulation studies, we estimated the parameter values and compared them to the values used to simulate the data. To assess the accuracy of the mass predictions, we calculated the root mean square error (RMSE) that compares the value of the predicted mass at time *t* for each individual *i*, 𝑚̂_𝑖,𝑡_, with the associated simulated mass value, 𝑚_𝑖,𝑡_. To ensure that the comparison is equivalent in the second simulation study, we did not include the estimated final mass in the RMSE, since it would only be available for the model without the mass at recapture. The code used for the simulations is available in the Supplementary Materials.

## Results

### Case study

The state-space model estimated the baseline mass loss as 4.68 g/hr (𝛽_0_ = 0.013 g/10 sec), with additional loss of 12.96 g/hr (𝛽_f_ = 0.036 g/10 sec) while flying (Table 1). The diving function indicated that relative mass gain while diving, when the covariates were equal to 0, was 25.56 g/hr (𝛽_d_ = 0.071 g/10 sec). As expected, mass gain increased with maximum depth and increased pitch standard deviation (i.e. birds gained more mass when making deeper dives with more changes in vertical direction). Re-fitting the model with the final mass excluded resulted in an RMSE in the estimated final mass of 33.7 g.

The state-space model enables us to examine detailed foraging success within a deployment; to illustrate this, we will focus on the mass estimates from a single individual (Figures 1-2). This murre was first captured at 10:35 UTC and recaptured for logger retrieval almost 24 hr later (2023-07-08 10:14 UTC). Starting and ending mass from the deployment was 1,084 and 1,082 g, respectively. When the bird departed on its foraging trip at 22:26 UTC, we estimated the mass to have declined to 1,026.9 g. Over a 9.3 hr foraging trip, the murre was estimated to have a net mass gain of 67.7 g. During the trip, estimated mass peaked at 1,108.0 g at 6:30 UTC and the bird began commuting back to the colony shortly after at 7:10 UTC. Most foraging occurred to the northwest of the colony. After reaching the maximum distance from the colony, the murre alternated bouts of foraging and resting on the water as it drifted east back toward the colony.

**Figure 1.**
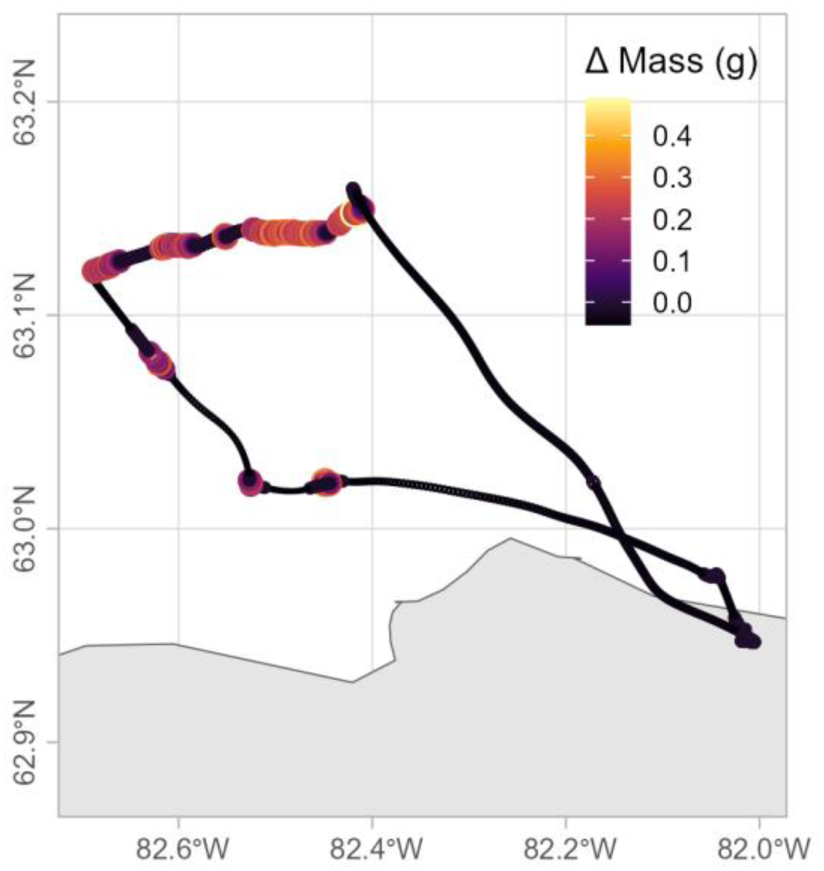
Example map showing estimated mass changes for a single deployment on an incubating thick-billed murre from Coats Island, NU. Colours and point sizes indicate estimated change in mass (g) at each time step, with larger points indicating higher mass changes.

**Figure 2.**
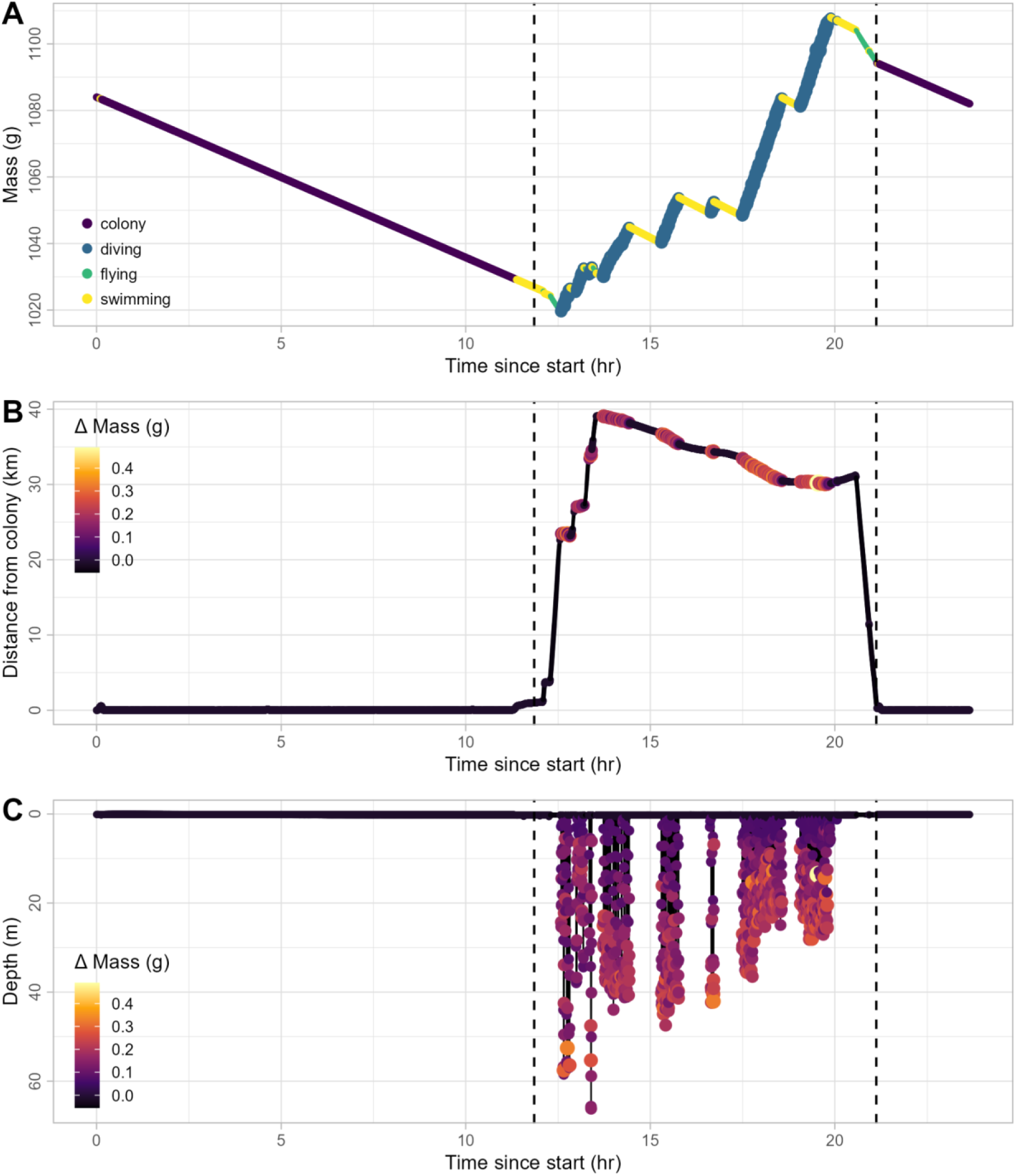
Example showing estimated mass changes for a single deployment on an incubating thick-billed murre from Coats Island, NU. Panel A shows the cumulative change in mass through time, colours indicate behaviour at each time step and the vertical lines indicate start and end of the foraging trip. Panel B shows distance from colony through time, colours and point sizes indicate estimated change in mass (g) at each time step, with larger points indicating higher mass changes. Panel C shows depth through time with colour and size scales as in B.

We examined estimated mass gain across all foraging trips to explore factors influencing mass balance and foraging success at the population level. Gross mass gain for foraging trips was 81.8 ± 34.5 g (range: 12.8-159.9 g) and net mass gain was 32.0 ± 31.9 g (range: -40.8-99.3 g, Figure 3). Based on the baseline mass loss from the model (𝛽_0_), a regular 12 hr shift at the colony would be associated with a loss of 57 g, therefore, murres should return from foraging with a net gain of at least 57 g to maintain their mass balance. Only 20.8% of foraging trips exceeded this threshold, suggesting that murres are incurring a mass deficit during incubation. There was no statistically significant relationship between trip duration and net mass gain (F_1, 46_ = 0.12, p = 0.73, Figure 4A), indicating that time away from the colony was not associated with mass balance. Net mass gain increased with maximum distance from the colony at a rate of 0.6 ± 0.3 g/km (F_1, 46_ = 5.01, p = 0.03, Figure 4B), suggesting that foraging success increased with distance from the colony. There was a weak, and not statistically significant, negative trend between starting mass and net mass gain (F_1, 46_ = 3.55, p = 0.07, Figure 4C).

**Figure 3.**
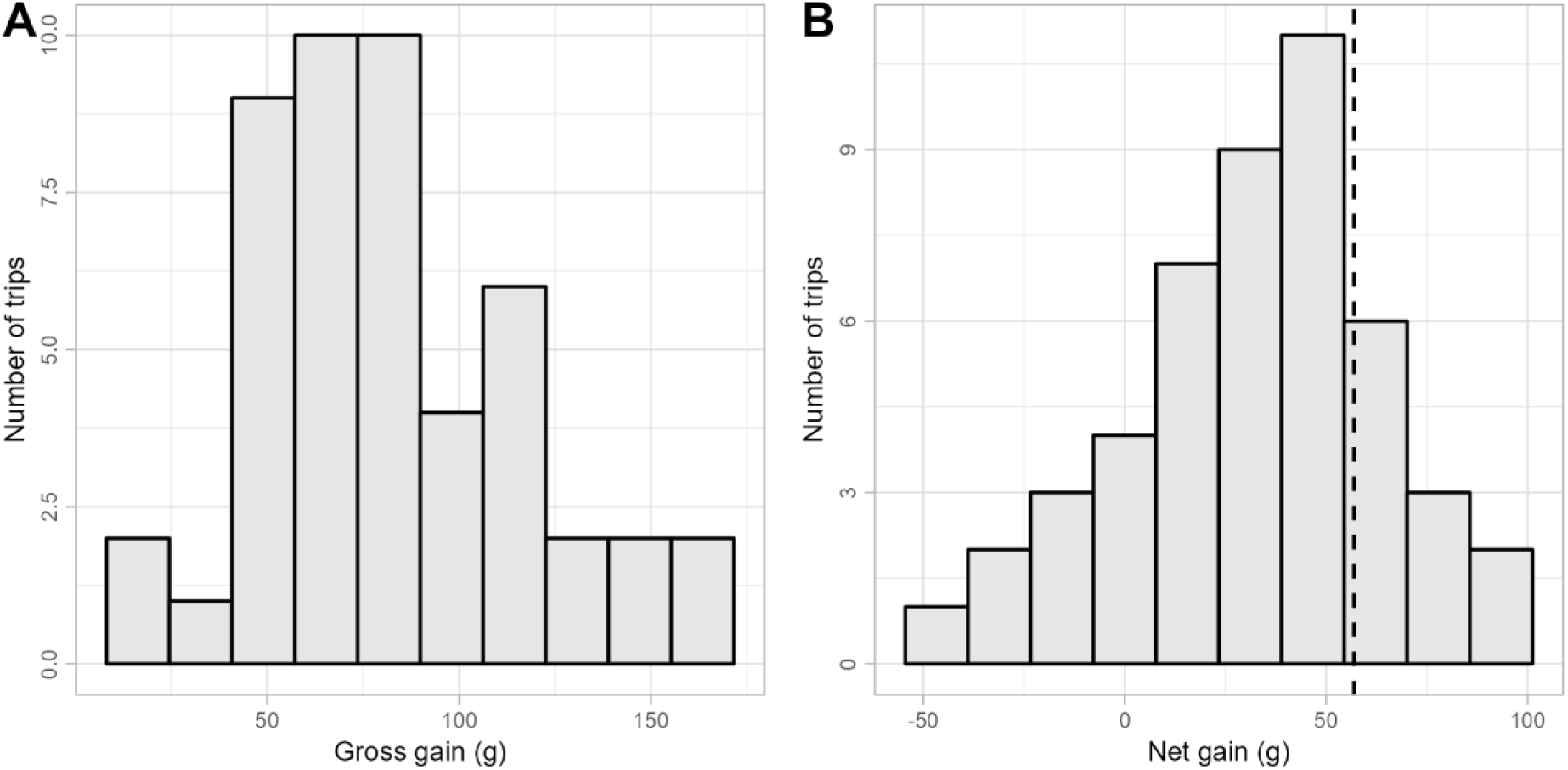
Histograms showing the distribution of (A) gross mass gain and (B) net mass gain across all foraging trips by incubating murres from Coats Island, NU. Dashed vertical line in panel B shows the estimated net mass gain needed to support a 12 hr shift of incubation.

**Figure 4.**
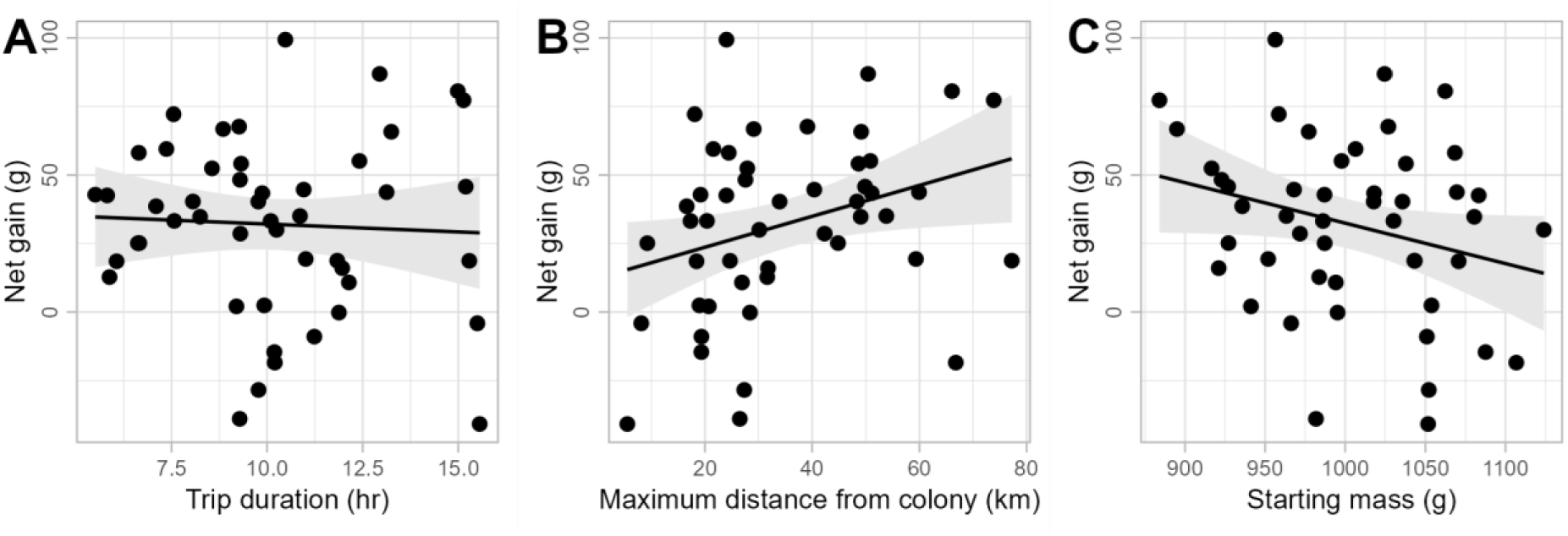
Effects of (A) trip duration, (B) maximum distance from colony, and (C) starting mass on net mass gain during foraging trips by incubating murres from Coats Island, NU. Lines show regression fit and shaded areas are 95% confidence intervals.

We examined the spatial distribution of gross gain rate while diving to identify regions with higher foraging potential. Gross mass gain generally increased with distance from the colony (Figure 5). The highest gain rates occurred to the northwest, north, and east of the colony. Uncertainty was higher to the north and east of the colony, because relatively few murres used these areas for foraging.

**Figure 5.**
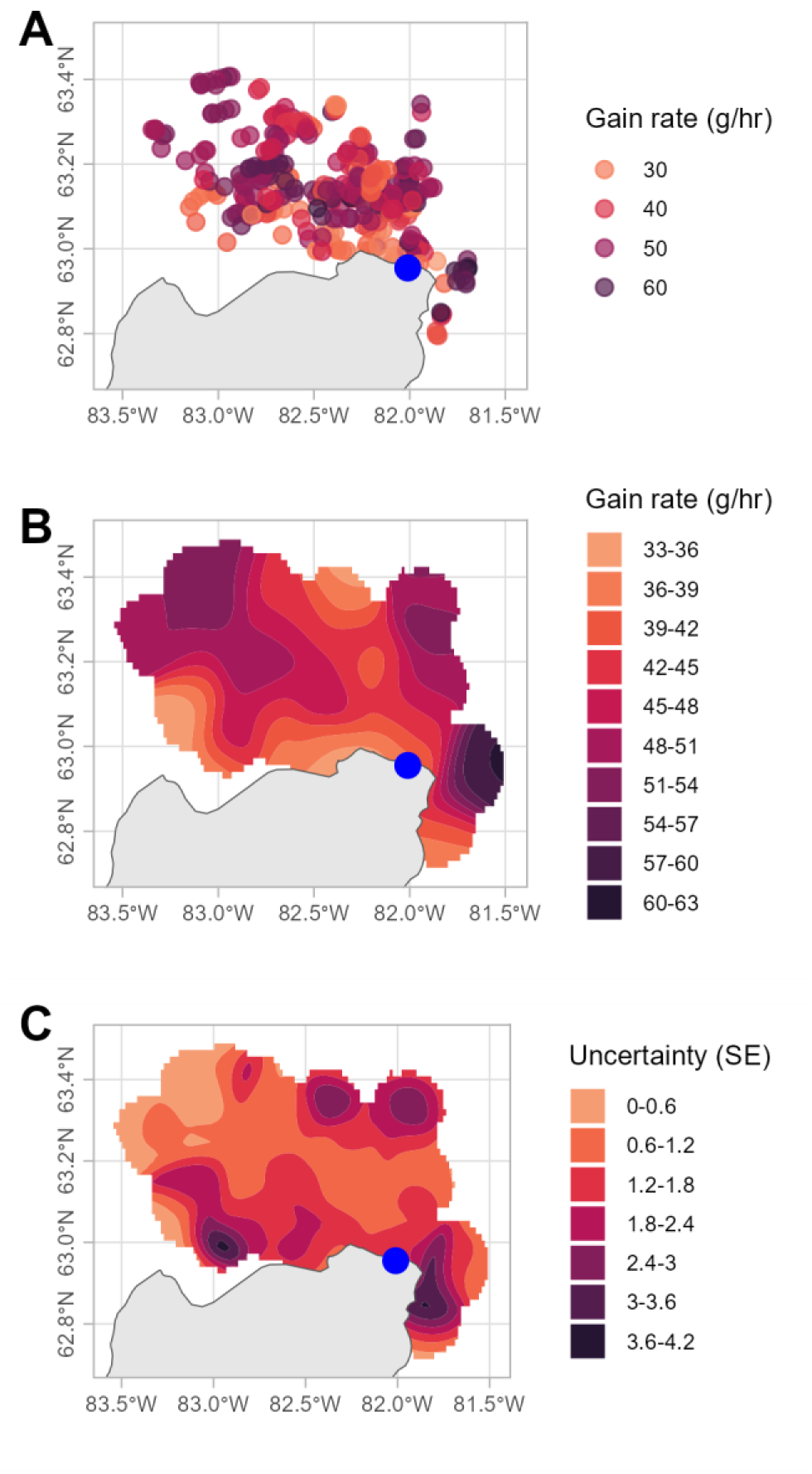
Spatial distribution of gain rate while diving for incubating thick-billed murres from Coats Island, NU. Panel A shows the hourly gain rates recorded from individual tracks. Panel B shows spatially smoothed average gain rate. Panel C shows the uncertainty in spatial estimates. Blue point indicates the location of the breeding colony.

### Simulation studies

The first simulation study demonstrated that, while the accuracy of the mass predictions decreased when the standard deviation of the process equation was misspecified, the main factor affecting the accuracy was the size of the true variation in mass (Figure 6A). Similarly, misspecification in the standard deviation of the process equation did not appear to be the main factor affecting the accuracy of the parameter estimates (Supplementary Materials A: Figure A1).

**Figure 6.**
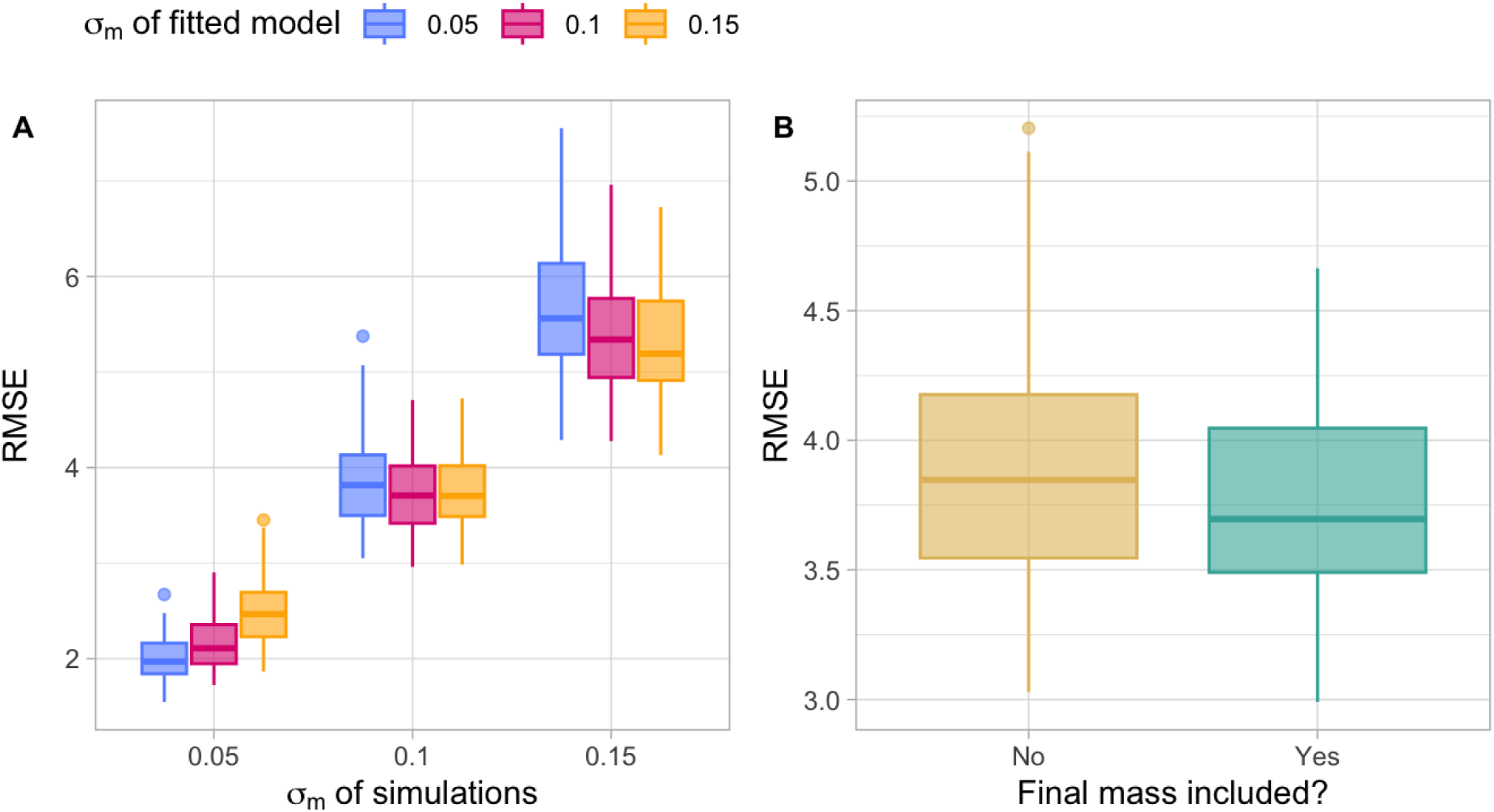
Effects of (A) the misspecification of the standard deviation that is associated with mass gain (𝜎_𝑚_) and (B) the inclusion of the final mass in the model on the root mean square error (RMSE). Panel (A) shows the results of the simulation study that modified both the standard deviation of the process equation in the simulations and the values fixed in the fitted model. Panel (B) shows the results of the simulation study fitted the same model with and without the final mass included as a fixed value in the model. Both panels only include the models that converged.

The results of the second simulation study indicated that including the final mass provided a small improvement in the accuracy of the mass predictions (Figure 6B), but doing so did not appear to noticeably improve the parameter estimates (Supplementary Materials A: Figure A2). Both simulation studies indicated that the estimates of the standard deviations of the random effect, 𝜎_𝑎_, and the observation equation, 𝜎_𝑤_, were biased, with estimates being consistently lower than the simulated values (Supplementary Materials A: Figures A1-A2). However, the other parameter estimates were generally close to the values used in the simulations. We note that 23% of the simulations (simulation study 1: 22.6%; simulation study 2: 27.5%) had convergence problems. We only considered the results from the converged models.

## Discussion

Our state-space model, based on simple biomechanical theory, estimated mass change and foraging success in birds using measures of wingbeat frequency. Our model could be applied to any animal engaging in flapping flight with identifiable foraging periods. We are in a “Golden Age of Biologging” and the location of millions of animals are now recorded (Kays et al., 2022; Wilmers et al., 2015). By recording acceleration, we show that mass gained can be measured providing insight into why they are where they are. Our case study also illustrates the utility of this approach for testing foraging theory and identifying critical foraging habitats. We provided simple examples of how results from the state-space model can be used to examine foraging success over multiple spatiotemporal scales. Our case study demonstrated that: dive profiles (maximum depth and dive complexity) influenced mass gained while foraging; most incubating murres were experiencing mass loss over a 24 hour foraging cycle; foraging trip success increased with distance from colony; and, gross mass gain was higher farther from the colony. Our method extends the novel approach of Sato et al. (2008), which estimated change in mass from changes in wingbeat frequencies between discrete bouts of flying, to provide a flexible new approach to model continuous mass change in flying species.

Field and captive studies support the relationship between mass and wingbeat frequency that drives our model. European shags had higher wingbeat frequency measured with accelerometers during incoming than outgoing flights (Sato et al., 2008; Watanuki et al., 2023). Changes in mass and wingbeat frequency of migrating Brent Geese (*Branta bernicla*) were consistent with Pennycuick’s predictions (Gudmundsson et al., 1995; Hedenström & Alerstam, 1992; Pennycuick, 1996). A wind tunnel study of Barn Swallows (*Hirundo rustica*) concluded that Pennycuick’s proposed scaling of wingbeat frequency with body mass applied at the individual level (Schmidt-Wellenburg et al., 2007); however, they observed a scaling exponent of 0.36 (95% CI 0.12-0.64), which was lower than, but overlapped, the proposed 0.5 scaling (Pennycuick, 1996). Similarly, observed metabolic power required to support increased mass is lower than predicted by theory (Kvist et al., 2001) Our state-space model could be readily modified to estimate the scaling exponent between mass and wingbeat frequency directly from the data.

We used a simple process equation to represent how murres lose and gain mass over time. Our case study demonstrated how this implementation can easily accommodate covariates for mass gain while foraging. In the future, the equation could be extended to provide a more realistic representation of this process. Using two separate equations for mass loss while fasting and mass gain while foraging would allow for differences in the variability of these two components’ energetics. Mass loss over time while fasting would have less variability within and among individuals than mass gained while foraging, which could be accommodated better with separate error terms. In the future, we would like to extend the model to include chick-feeding events for birds that are attending chicks. This would allow us to directly estimate energetic investment in the chick relative to the self; this would also provide a test of model accuracy in cases where the size of chick meals can be independently measured through feeding observations.

There was significant variation in observed wingbeat frequency (*w*) values; future work could adapt the observation equation to better account for variables that influence *w,* such as flight manoeuvres, air density, turbulence, wind speed and direction, and wingbeat amplitude. The relationship between *m* and *w* is for birds in constant level flight, excluding periods when birds are engaged in flight manoeuvres (e.g. take-off/landing, changing altitude, turning) could reduce noise in the observation equation. Altered wingbeat kinematics will be needed to power ascent (Van Walsum et al., 2020), and identifying periods of level constant flight will be helpful. Wingbeat frequency should increase when air density decreases in order to compensate for decreasing lift (Pennycuick, 1996; Schmaljohann & Liechti, 2009; Shepard et al., 2024). Air density changes with temperature and altitude, so it may be important to account for air density for species that fly at different heights and in complex landscapes. However, seabirds flying at sea level over the ocean may not experience significant fluctuation in air density over short term studies like ours. Wingbeat frequency increases with airspeed (which will be higher flying into the wind than with a tailwind), which is a biomechanical necessity to keep the bird flying faster (Elliott et al., 2014; Kogure et al., 2016). Although it may be challenging to obtain measures of wind dynamics at a spatial and temporal scale relevant to bird movements, accounting for wind speed and direction could account for unexplained variability in wingbeat frequency. Finally, Sato et al. (2008) note that mass change should be proportional to wingbeat amplitude times frequency; therefore, incorporating wingbeat amplitude, as well as other components of kinematics (i.e. upstroke to downstroke ratio), could be helpful in refining the observation equation to isolate the relationship between wingbeat frequency and mass. Theoretically, a similar biomechanical approach could be used for wing-propelled divers, using changes in stroke-frequency during descents of known body angle to estimate changes in mass, if airspace volume were estimated from variation in ascent rate, which could be incorporated into the model as a second observation equation.

Currently, the model focuses on discrete foraging periods - diving birds in our case study - with flapping flight. It could likely be expanded to flap-gliding birds and non-diving species. However, behavioural identification of foraging periods would be necessary to include covariates on the mass gain component of the model. State-space models can be hard to fit to data, and in particular, it can be difficult to simultaneously estimate the parameters associated with biological stochasticity and measurement error (Auger-Méthé et al., 2016, 2021). The case study and the simulation studies demonstrated that while our state-space model is not impervious to such problems, some modelling decisions can help the estimation. First, we ran into convergence problems, highlighting the importance of checking convergence. We resolved the convergence issue in the case study by fixing the standard deviation parameter associated with the biological stochasticity (𝜎_𝑚_). The simulation study showed that the decrease in the accuracy of the mass estimates associated with misspecifying this parameter was relatively small, suggesting that as long as a biologically relevant 𝜎_𝑚_ is chosen, the bias in mass predictions should be relatively small. Second, while we built the model to always include the mass of the bird at capture, we also recommend including the mass at recapture. Fitting the model to our case study without the final mass resulted in an RMSE in the estimated final mass of 33.7 g. The simulation study showed an improvement in the accuracy of the mass estimates when the final mass was included. While the accuracy gains were relatively small, we expect that in most real applications, in which the model rarely fully captures the underlying mechanism, including the final mass will significantly improve the accuracy of the mass predictions. The size of the difference between the final mass predicted for the murre case study when mass at recapture was ignored and the measured mass further suggests that including mass at recapture is highly beneficial. Finally, the simulation study indicated that the estimated standard deviation for the individual random effect and the measurement error were negatively biased. We note that due to the computational cost of running these simulations and their dependence on the limited amount of murre data, we simulated 20 tracks. A preliminary study comparing simulations with 10 individual tracks to those obtained with 20 showed that increasing the number of tracks improves the model fit. While further increasing the sample would likely diminish the magnitude of the bias, we note that estimating multiple levels of stochasticity is a known challenge of state-space models (Auger-Méthé et al., 2016, 2021). We emphasize that the simulation studies show that other parameter estimates and the predicted mass were unbiased and that these reliable estimates and predictions are generally of greater importance in answering ecological questions.

Our method can answer key questions in wildlife ecology and conservation. For example, the mass gained at sea can be used to create energyscapes and define hotspots for marine protected areas, assuming that areas of consistent high energy intake are critical habitats. Moreover, linking foraging behaviour with fitness has been a longstanding challenge in movement ecology, likely because we seldom have direct measures of foraging success. Because of their small size and low power consumption allowing study of foraging behaviour in a range of species and over long durations, there are now long-term datasets on accelerometers (i.e. 15 years at our study site). Variation among years in foraging success should, in principle, be closely related to fitness, improving mechanistic models linking environment to fitness. Individual foraging strategies could be related to mass gained at sea, allowing us to examine why different strategies exist. Our simple model, using data for miniature accelerometers that are widely used in wildlife ecology, should open the door to answering many key questions that depend on one of ecology’s most important parameters, energy intake.

## Acknowledgements

HG Gilchrist, H Hennin, A Eby, M Gousy-LeBlanc, F Tremblay, D Noblet, and J Angootealuk assisted with data collection for the case study. Field research was undertaken with funding from the Natural Sciences and Engineering Research Council (NSERC) of Canada, Environment and Climate Change Canada, the Northern Contaminants Program, and the Newfoundland and Labrador Murre Fund with support from Wildlife Habitat Canada and Bird Studies Canada. AP received support from the Weston Family Foundation. MAM thanks the Natural Sciences and Engineering Research Council of Canada (NSERC), the Canadian Research Chairs program, BC Knowledge Development fund and Canada Foundation for Innovation’s John R. Evans Leaders Fund for their financial support.

## Conflict of interest

The authors have no conflicts of interest to report.

## Author contributions

All authors conceived the ideas and designed methodology; AP and KE collected the data; AP and MAM analysed the data. All authors contributed to writing the manuscript, revised the drafts and gave final approval for publication.

## Animal care permits

All animal handling was conducted under permit McGill University Animal Use permit 7599.

## Supplementary Materials A: Additional simulation results

**Figure A1.**
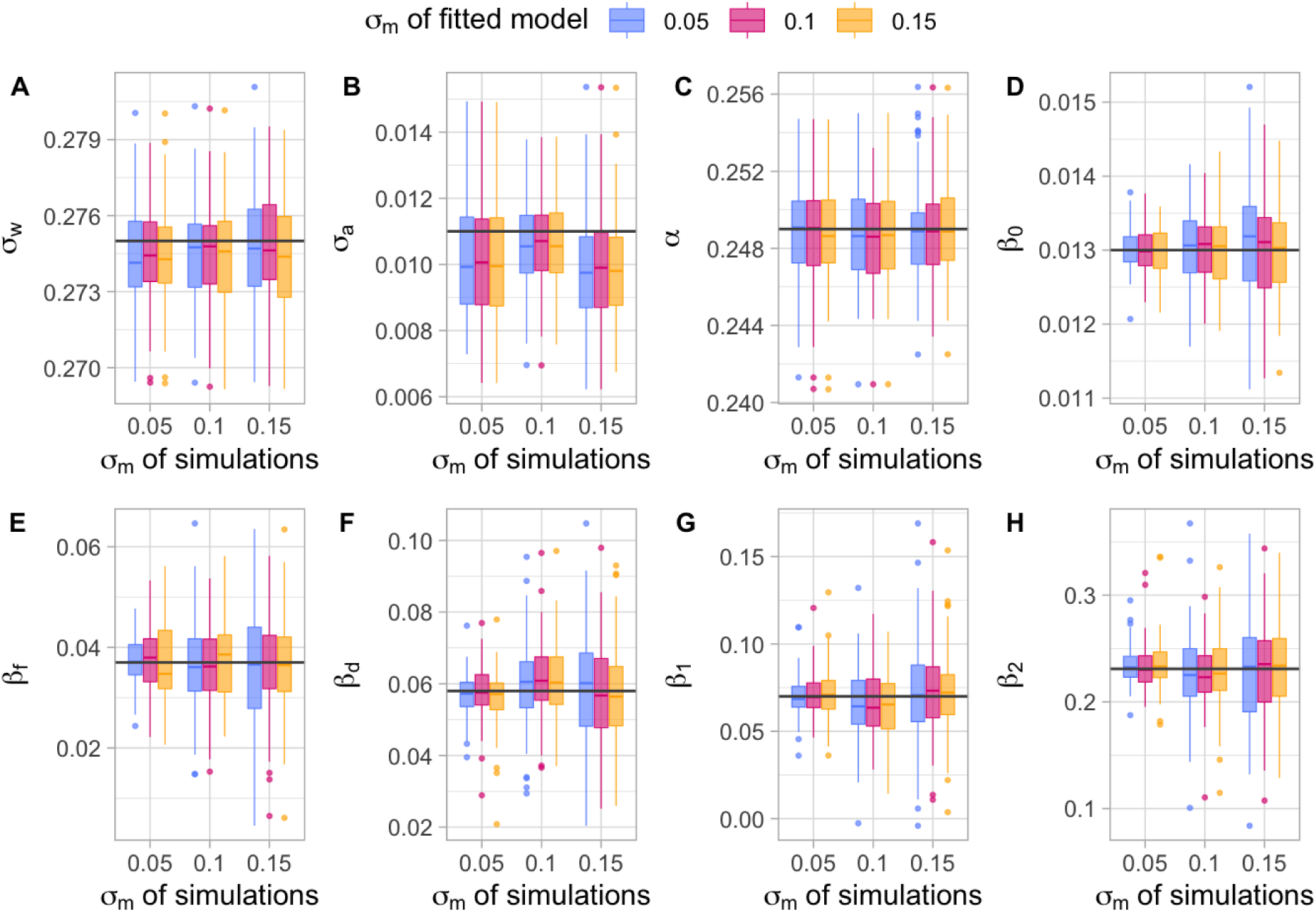
Effects of the misspecification of the standard deviation that is associated with mass gain (𝜎_𝑚_) on the parameter estimates. Each panel shows the parameter estimate values for simulations that varied in terms of the standard deviation of the process equation used in both the simulations and the values fixed in the fitted model. All panels only include the models that converged. The grey lines represent the value of the parameter used in the simulations.

**Figure A2.**
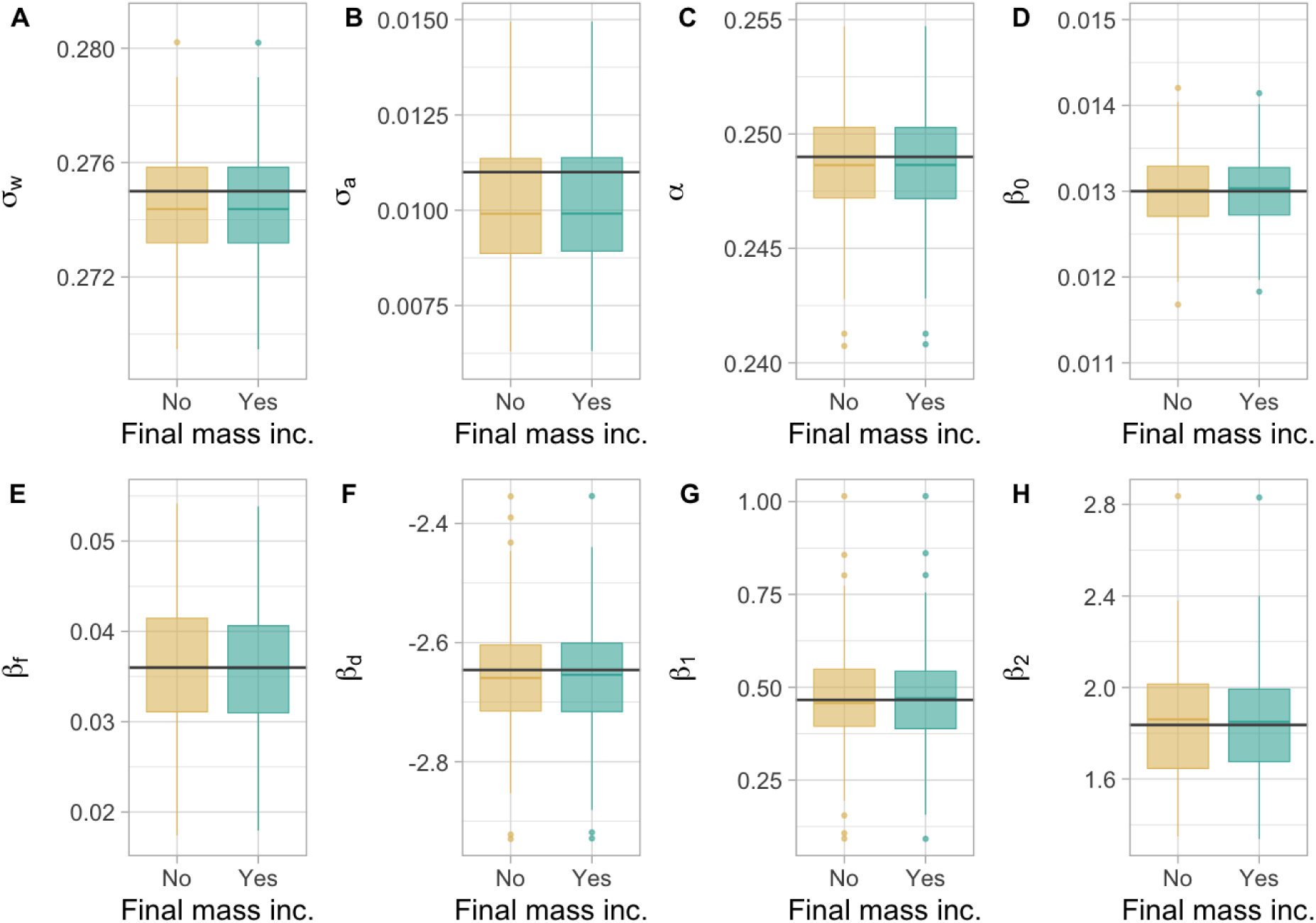
Effects of the inclusion of the final mass in the model on the parameter estimates. Each panel shows the parameter estimate values for simulations fitted the same model as the one simulated but that varied in whether the final mass was included as a fixed value in the model. All panels only include the models that converged. The grey lines represent the value of the parameter used in the simulations.

## Notes

### Competing Interest Statement

The authors have declared no competing interest.

## References

1. Auger-Méthé, M., Field, C., Albertsen, C. M., Derocher, A. E., Lewis, M. A., Jonsen, I. D., & Mills Flemming, J. (2016). State-space models’ dirty little secrets: Even simple linear Gaussian models can have estimation problems. Scientific Reports, 6(1), 26677. 10.1038/srep26677

2. Auger-Méthé, M., Newman, K., Cole, D., Empacher, F., Gryba, R., King, A. A., Leos-Barajas, V., Mills Flemming, J., Nielsen, A., Petris, G., & Thomas, L. (2021). A guide to state–space modeling of ecological time series. Ecological Monographs, 91(4), e01470. 10.1002/ecm.1470

3. Baddeley, A., Rubak, E., & Turner, R. (2016). Spatial point patterns: Methodology and applications with R (Vol. 1). CRC Press.

4. Bennison, A., Bearhop, S., Bodey, T. W., Votier, S. C., Grecian, W. J., Wakefield, E. D., Hamer, K. C., & Jessopp, M. (2018). Search and foraging behaviors from movement data: A comparison of methods. Ecology and Evolution, 8(1), 13–24. 10.1002/ece3.3593

5. Biuw, M., Boehme, L., Guinet, C., Hindell, M., Costa, D., Charrassin, J.-B., Roquet, F., Bailleul, F., Meredith, M., Thorpe, S., Tremblay, Y., McDonald, B., Park, Y.-H., Rintoul, S. R., Bindoff, N., Goebel, M., Crocker, D., Lovell, P., Nicholson, J., … Fedak, M. A. (2007). Variations in behavior and condition of a Southern Ocean top predator in relation to in situ oceanographic conditions. Proceedings of the National Academy of Sciences, 104(34), 13705–13710. 10.1073/pnas.0701121104

6. Biuw, M., McConnell, B., Bradshaw, C. J. A., Burton, H., & Fedak, M. (2003). Blubber and buoyancy: Monitoring the body condition of free-ranging seals using simple dive characteristics. Journal of Experimental Biology, 206(19), 3405–3423. 10.1242/jeb.00583

7. Brisson-Curadeau, É., Handrich, Y., Elliott, K. H., & Bost, C.-A. (2021). Accelerometry predicts prey-capture rates in the deep-diving king penguin Aptenodytes patagonicus. Marine Biology, 168(10), 156. 10.1007/s00227-021-03968-y

8. Butler, P. J., Green, J. A., Boyd, I. L., & Speakman, J. R. (2004). Measuring metabolic rate in the field: The pros and cons of the doubly labelled water and heart rate methods. Functional Ecology, 18(2), 168–183. 10.1111/j.0269-8463.2004.00821.x

9. Charnov, E. L. (1976). Optimal foraging, the marginal value theorem. Theoretical Population Biology, 9(2), 129–136. 10.1016/0040-5809(76)90040-X

10. Chimienti, M., Cornulier, T., Owen, E., Bolton, M., Davies, I. M., Travis, J. M. J., & Scott, B. E. (2016). The use of an unsupervised learning approach for characterizing latent behaviors in accelerometer data. Ecology and Evolution, 6(3), 727–741. 10.1002/ece3.1914

11. Clermont, J., Woodward-Gagné, S., & Berteaux, D. (2021). Digging into the behaviour of an active hunting predator: Arctic fox prey caching events revealed by accelerometry. Movement Ecology, 9(1), 58. 10.1186/s40462-021-00295-1

12. Del Raye, G., Jorgensen, S. J., Krumhansl, K., Ezcurra, J. M., & Block, B. A. (2013). Travelling light: White sharks (Carcharodon carcharias) rely on body lipid stores to power ocean-basin scale migration. Proceedings of the Royal Society B: Biological Sciences, 280(1766), 20130836. 10.1098/rspb.2013.0836

13. Elliott, K. H., Chivers, L. S., Bessey, L., Gaston, A. J., Hatch, S. A., Kato, A., Osborne, O., Ropert-Coudert, Y., Speakman, J. R., & Hare, J. F. (2014). Windscapes shape seabird instantaneous energy costs but adult behavior buffers impact on offspring. Movement Ecology, 2(1), 17. 10.1186/s40462-014-0017-2

14. Elliott, K. H., Le Vaillant, M., Kato, A., Speakman, J. R., & Ropert-Coudert, Y. (2013). Accelerometry predicts daily energy expenditure in a bird with high activity levels. Biology Letters, 9(1), 20120919. 10.1098/rsbl.2012.0919

15. Elliott, K. H., Ricklefs, R. E., Gaston, A. J., Hatch, S. A., Speakman, J. R., & Davoren, G. K. (2013). High flight costs, but low dive costs, in auks support the biomechanical hypothesis for flightlessness in penguins. Proceedings of the National Academy of Sciences, 110(23), 9380– 9384. 10.1073/pnas.1304838110

16. Florko, K. R. N., Shuert, C. R., Cheung, W. W. L., Ferguson, S. H., Jonsen, I. D., Rosen, D. A. S., Sumaila, U. R., Tai, T. C., Yurkowski, D. J., & Auger-Méthé, M. (2023). Linking movement and dive data to prey distribution models: New insights in foraging behaviour and potential pitfalls of movement analyses. Movement Ecology, 11(1), 17. 10.1186/s40462-023-00377-2

17. Green, J. A. (2011). The heart rate method for estimating metabolic rate: Review and recommendations. Comparative Biochemistry and Physiology Part A: Molecular & Integrative Physiology, 158(3), 287–304. 10.1016/j.cbpa.2010.09.011

18. Gudmundsson, G. A., Benvenuti, S., Alerstam, T., Papi, F., Lilliendahl, K., & Åkesson, S. (1995). Examining the Limits of Flight and Orientation Performance: Satellite Tracking of Brent Geese Migrating across the Greenland Ice-Cap. Proceedings: Biological Sciences, 261(1360), 73–79.

19. Hambly, C., Harper, E. J., & Speakman, J. R. (2004). The energy cost of loaded flight is substantially lower than expected due to alterations in flight kinematics. Journal of Experimental Biology, 207(22), 3969–3976. 10.1242/jeb.01234

20. Hedenström, A., & Alerstam, T. (1992). Climbing Performance of Migrating Birds as A Basis for Estimating Limits for Fuel-Carrying Capacity and Muscle Work. Journal of Experimental Biology, 164(1), 19–38. 10.1242/jeb.164.1.19

21. Hicks, O., Burthe, S., Daunt, F., Butler, A., Bishop, C., & Green, J. A. (2017). Validating accelerometry estimates of energy expenditure across behaviours using heart rate data in a free-living seabird. Journal of Experimental Biology, 220(10), 1875–1881. 10.1242/jeb.152710

22. Jonsen, I. D., Grecian, W. J., Phillips, L., Carroll, G., McMahon, C., Harcourt, R. G., Hindell, M. A., & Patterson, T. A. (2023). aniMotum, an R package for animal movement data: Rapid quality control, behavioural estimation and simulation. Methods in Ecology and Evolution, 14(3), 806–816. 10.1111/2041-210X.14060

23. Kays, R., Davidson, S. C., Berger, M., Bohrer, G., Fiedler, W., Flack, A., Hirt, J., Hahn, C., Gauggel, D., Russell, B., Kölzsch, A., Lohr, A., Partecke, J., Quetting, M., Safi, K., Scharf, A., Schneider, G., Lang, I., Schaeuffelhut, F., … Wikelski, M. (2022). The Movebank system for studying global animal movement and demography. Methods in Ecology and Evolution, 13(2), 419–431. 10.1111/2041-210X.13767

24. Kogure, Y., Sato, K., Watanuki, Y., Wanless, S., & Daunt, F. (2016). European shags optimize their flight behavior according to wind conditions. Journal of Experimental Biology.

25. Kooyman, G. L., & Ponganis, P. J. (1998). The physiological basis of diving to depth: Birds and mammals. Annual Review of Physiology, 60, 19–32. 10.1146/annurev.physiol.60.1.19

26. Kristensen, K., Nielsen, A., Berg, C. W., Skaug, H., & Bell, B. M. (2016). TMB: Automatic Differentiation and Laplace Approximation. Journal of Statistical Software, 70, 1–21. 10.18637/jss.v070.i05

27. Kvist, A., Lindström, Å., Green, M., Piersma, T., & Visser, G. H. (2001). Carrying large fuel loads during sustained bird flight is cheaper than expected. Nature, 413(6857), 730–732. 10.1038/35099556

28. Lok, T., van der Geest, M., Bom, R. A., de Goeij, P., Piersma, T., & Bouten, W. (2023). Prey ingestion rates revealed by back-mounted accelerometers in Eurasian spoonbills. Animal Biotelemetry, 11(1), 5. 10.1186/s40317-022-00315-w

29. Lovvorn, J. R., Watanuki, Y., Kato, A., Naito, Y., & Liggins, G. A. (2004). Stroke patterns and regulation of swim speed and energy cost in free-ranging Brünnich’s guillemots. Journal of Experimental Biology, 207(26), 4679–4695. 10.1242/jeb.01331

30. MacArthur, R. H., & Pianka, E. R. (1966). On Optimal Use of a Patchy Environment. The American Naturalist, 100(916), 603–609.

31. Orians, G., & Pearson, N. (1979). On the theory of central place foraging. In Analysis of ecological systems (pp. 155–177). Ohio State Press.

32. Patterson, A., Gilchrist, H. G., Chivers, L., Hatch, S., & Elliott, K. (2019). A comparison of techniques for classifying behavior from accelerometers for two species of seabird. Ecology and Evolution, 9(6), 3030–3045. 10.1002/ece3.4740

33. Pennycuick, C. J. (1996). Wingbeat Frequency of Birds in Steady Cruising Flight: New Data and Improved Predictions. Journal of Experimental Biology, 199(7), 1613–1618. 10.1242/jeb.199.7.1613

34. R Core Team. (2023). R: A language and environment for statistical computing (Version 4.3.1) [Computer software]. R Foundation for Statistical Computing.

35. Robinson, P. W., Simmons, S. E., Crocker, D. E., & Costa, D. P. (2010). Measurements of foraging success in a highly pelagic marine predator, the northern elephant seal. Journal of Animal Ecology, 79(6), 1146–1156. 10.1111/j.1365-2656.2010.01735.x

36. Ropert-Coudert, Y., & Kato, A. (2006). Are stomach temperature recorders a useful tool for determining feeding activity? Polar Biosciences, 20, 63–72.

37. Saldanha, S., Cox, S. L., Militão, T., & González-Solís, J. (2023). Animal behaviour on the move: The use of auxiliary information and semi-supervision to improve behavioural inferences from Hidden Markov Models applied to GPS tracking datasets. Movement Ecology, 11(1), 41. 10.1186/s40462-023-00401-5

38. Sato, K., Daunt, F., Watanuki, Y., Takahashi, A., & Wanless, S. (2008). A new method to quantify prey acquisition in diving seabirds using wing stroke frequency. Journal of Experimental Biology, 211(1), 58–65. 10.1242/jeb.009811

39. Sato, K., Watanuki, Y., Takahashi, A., Miller, P. J. O., Tanaka, H., Kawabe, R., Ponganis, P. J., Handrich, Y., Akamatsu, T., Watanabe, Y., Mitani, Y., Costa, D. P., Bost, C.-A., Aoki, K., Amano, M., Trathan, P., Shapiro, A., & Naito, Y. (2007). Stroke frequency, but not swimming speed, is related to body size in free-ranging seabirds, pinnipeds and cetaceans. Proceedings of the Royal Society B: Biological Sciences, 274(1609), 471–477. 10.1098/rspb.2006.0005

40. Schick, R. S., New, L. F., Thomas, L., Costa, D. P., Hindell, M. A., McMahon, C. R., Robinson, P. W., Simmons, S. E., Thums, M., Harwood, J., & Clark, J. S. (2013). Estimating resource acquisition and at-sea body condition of a marine predator. Journal of Animal Ecology, 82(6), 1300–1315. 10.1111/1365-2656.12102

41. Schmaljohann, H., & Liechti, F. (2009). Adjustments of wingbeat frequency and air speed to air density in free-flying migratory birds. Journal of Experimental Biology, 212(22), 3633–3642. 10.1242/jeb.031435

42. Schmidt-Wellenburg, C. A., Biebach, H., Daan, S., & Visser, G. H. (2007). Energy expenditure and wing beat frequency in relation to body mass in free flying Barn Swallows (Hirundo rustica). Journal of Comparative Physiology B, 177(3), 327–337. 10.1007/s00360-006-0132-5

43. Shepard, E., Garde, B., Krishnan, K., Fell, A., Tatayah, V., Jones, C. G., Cole, N. C., & Lempidakis, E. (2024). Latitudinal gradients in air density create invisible topography at sea level, affecting animal flight costs. Current Biology, 34(24), 5846–5851.e4. 10.1016/j.cub.2024.10.064

44. Shepard, E., Wilson, R., Halsey, L., Quintana, F., Laich, A. G., Gleiss, A., Liebsch, N., Myers, A., & Norman, B. (2008). Derivation of body motion via appropriate smoothing of acceleration data. Aquatic Biology, 4(3), 235–241. 10.3354/ab00104

45. Shepard, E., Wilson, R., Quintana, F., Gómez Laich, A., Liebsch, N., Albareda, D., Halsey, L., Gleiss, A., Morgan, D., Myers, A., Newman, C., & McDonald, D. (2008). Identification of animal movement patterns using tri-axial accelerometry. Endangered Species Research, 10, 47–60. 10.3354/esr00084

46. Sidrow, E., Heckman, N., McRae, T. M., Volpov, B. L., Trites, A. W., Fortune, S. M. E., & Auger-Méthé, M. (2024). *Incorporating sparse labels into hidden Markov models using weighted likelihoods improves accuracy and interpretability in biologging studies* (arXiv:2409.18091). arXiv. 10.48550/arXiv.2409.18091

47. Stephens, D. W., & Krebs, J. R. (1986). Foraging theory (Vol. 6). Princeton university press.

48. Van Walsum, T. A., Perna, A., Bishop, C. M., Murn, C. P., Collins, P. M., Wilson, R. P., & Halsey, L. G. (2020). Exploring the relationship between flapping behaviour and accelerometer signal during ascending flight, and a new approach to calibration. Ibis, 162(1), 13–26. 10.1111/ibi.12710

49. Watanabe, Y. Y., & Takahashi, A. (2013). Linking animal-borne video to accelerometers reveals prey capture variability. Proceedings of the National Academy of Sciences, 110(6), 2199–2204. 10.1073/pnas.1216244110

50. Watanuki, Y., Sato, K., Shiomi, K., Wanless, S., & Daunt, F. (2023). Foraging habitat and site selection do not affect feeding rates in European shags. Journal of Experimental Biology, 226(4), jeb244461. 10.1242/jeb.244461

51. Wilmers, C. C., Nickel, B., Bryce, C. M., Smith, J. A., Wheat, R. E., & Yovovich, V. (2015). The golden age of bio-logging: How animal-borne sensors are advancing the frontiers of ecology. Ecology, 96(7), 1741–1753. 10.1890/14-1401.1

52. Wilson, R. P., Börger, L., Holton, M. D., Scantlebury, D. M., Gómez-Laich, A., Quintana, F., Rosell, F., Graf, P. M., Williams, H., Gunner, R., Hopkins, L., Marks, N., Geraldi, N. R., Duarte, C. M., Scott, R., Strano, M. S., Robotka, H., Eizaguirre, C., Fahlman, A., & Shepard, E. L. C. (2020). Estimates for energy expenditure in free-living animals using acceleration proxies: A reappraisal. Journal of Animal Ecology, 89(1), 161–172. 10.1111/1365-2656.13040

53. Wilson, R. P., Pütz, K., Grémillet, D., Culik, B. M., Kierspel, M., Regel, J., Bost, C. A., Lage, J., & Cooper, J. (1995). Reliability of Stomach Temperature Changes in Determining Feeding Characteristics of Seabirds. Journal of Experimental Biology, 198(5), 1115–1135. 10.1242/jeb.198.5.1115

54. Yong, C., Harcourt, R., McMahon, C. R., Costa, D. P., Huckstadt, L. A., Hindell, M., & Jonsen, I. (2024). Dive descent rate as a buoyancy indicator to infer body condition of Weddell seals in the Antarctic. Marine Mammal Science, 40(4), e13147. 10.1111/mms.13147

55. Zuur, A. F. (2007). Analysing Ecological Data. Statistics for Biology and Health/Springer.

