## Supplementary material for "Using wingbeat frequency to estimate mass gained by seabirds": case-study.html

Using wingbeat frequency to estimate mass gained by seabirds. Supplementary material B: case study


### Using wingbeat frequency to estimate mass gained by seabirds. Supplementary material B: case study

#### 2025-05-13

##### Compile C++ Models

Compile and load the c++ model named *wbf\_mass\_model.cpp*,
which contains the function to calculate the negative log likelihood of
the model.

```
# Load C++ model
compile("wbf_mass_model.cpp")
```

```
## [1] 0
```

```
dyn.load(dynlib("wbf_mass_model"))
```

##### Import data

Load the case study dataset, which comprise of the tracking data
collected from 55 logger deployments on thick-billed murres (*Uria
lomvia*) breeding at Coats Island, NU (82.01˚ W, 52.95˚ N) in 2023.
Biologgers were programmed to collect GPS (1 min), depth (1 Hz), and
tri-axial acceleration (50 Hz). Murres were recaptured 12-48 hours later
for device retrieval. Each murre was weighed to the closest gram at
initial deployment and at recapture. All biologger data were summarized
at 10-sec intervals for analysis. Variables in the data set are:

- bird\_id: Unique identifier for each deployment, string
- time: Timestamp for each observation, datetime
- wbf: Wing-beat frequencies (Hz), numeric
- flying: Indicator that bird is in flight, boolean
- diving: Indicator that bird is diving, boolean
- behaviour: Behavioural category (colony, diving, flying, swimming),
  factor
- depth\_m: Current depth (m), numeric
- max\_depth\_m: Maximum depth attained within a dive (m), numeric
- pitch\_sd: Standard deviation in pitch, numeric
- coldist: Distance from the colony (km), numeric
- lon: Longitude (decimal degrees), numeric
- lat: Latitude (decimal degrees), numeric
- mass\_on: Bird mass at start of deployment (g), numeric
- mass\_off: Bird mass at end of deployment (g), numeric
- trip\_id: Identifier for trips within deployments, numeric

```
dat <- read.csv('case_study_data.csv')
dat$time <- as.POSIXct(dat$time, format = '%Y-%m-%d %H:%M:%S', tz = 'UTC')
```

We make a small matrix that indexes the start and end row for each
individual in the dataset. This indexing will allow the C++ code to
differentiate among deployments as it fits the model. C++ uses indexing
that starts at 0, unlike R, which starts indexing at 1. It is important
to be aware of this difference as we prepare the data for the model.

```
# make row index that starts at 0 
dat$idx <- 0:(nrow(dat) - 1)

# summary of bird start and end rows using C++ numbering
bid <- dat |> 
  group_by(bird_id) |> 
  summarize(
    start = min(idx),
    end = max(idx)
  )

head(bid)
```

```
## # A tibble: 6 × 3
##   bird_id start   end
##     <int> <int> <int>
## 1       0     0  2526
## 2       1  2527  8401
## 3       2  8402 13952
## 4       3 13953 21999
## 5       4 22000 31648
## 6       5 31649 37307
```

The TMB model requires starting values for mass estimates. Below we
use our matrix of row indices [bid] to extract the initial mass [rdm0]
and final mass [rdmT] for each bird. Then we add a variable named
*mdata* to our dataframe, which will hold the the starting values
for estimating mass at each time step. We use the initial mass as the
starting value for all mass estimates for each bird, which will ensure
that our starting values are biologically reasonable.

The TMB model also requires a map indicating the mass values to be
estimated and those that are fixed. We already know the initial and
final mass, so we will map these observations to NA, which tells TMB to
fix these values. All other mass values are mapped with values from
`1:nrow(dat)`. In our case, it is important that all mapped
values are unique. If there are any repeating values in the mapping, TMB
will force masses with the same mapped values to be the same.

```
# Vectors of start and end mass by bird
rdm0 <- dat$mass_on[bid$start + 1]
rdmT <- dat$mass_off[bid$end + 1]

# Starting values for mass estimates
dat$mdata <- NA
dat$mdata[(bid$start) + 1] <- rdm0
dat$mdata[(bid$end) + 1] <- rdmT
dat$mdata <- imputeTS::na_locf(dat$mdata) # fill with starting mass
```

```
## Registered S3 method overwritten by 'quantmod':
##   method            from
##   as.zoo.data.frame zoo
```

```
# Fix start and end mass to input values
dat$mmap <- 1:nrow(dat) # make sure all map values to be estimate are unique
dat$mmap[bid$start + 1] <- NA # fix start values to known starting mass
dat$mmap[bid$end + 1] <- NA # fix end values to known ending mass

# Convert to a matrix for TMB
bid <- as.matrix(bid)
```

##### Run model

Here we prepare our inputs for TMB. We set up the data inputs as a
named list with all the data expected by the TMB model:

- w: a vector of wingbeat frequency values
- d: a vector of 0/1 indicating when birds are diving
- f: a vector of 0/1 indicating when birds are flying
- c1: a vector of the first covariate for the mass gain equation
- c2: a vector of the second covariate for the mass gain equation
- bid: a matrix indexing the start and end rows for each bird using
  C++ indexing

Note that both c1 and c2 have been scaled so that values range from
0-1 before being based to dataTMB.

The c++ code be modified to change (add or remove) the number of
covariates influencing mass gain while foraging. In which case, you
would change the number of covariates passed to dataTMB.

```
# Set a fixed value for variance in the process equation
sdp <- 0.1

# Data object to pass to TMB
dataTmb <- list(w = dat$wbf, # Vector of WBF values, includes NAs when not in flight
                d = dat$diving, # Vector of 0/1 indicating when birds are diving/foraging
                f = dat$flying, # Vector of 0/1 indicating when birds are flying
                c1 = (dat$max_depth_m)/max(dat$max_depth_m), # Covariate for maximum dive depth, scaled to take values between 0-1
                c2 = (dat$pitch_sd)/max(dat$pitch_sd), # Covariate for SD in pitch, scaled to take values between 0-1
                bid = bid # Matrix of row indicators for each bird
)
```

Next, we create a named list of the starting values we will pass to
TMB. This list includes all estimated and fixed parameters used in the
model. We have included 9 coefficients that are part of the process and
observation equations. There is also a vector `ai` the same
length as the number of deployments in data and the vector for our
starting mass values `dat$mdata`.

Certain parameters should only take values > 0. These are passed
to TMB on the log scale and exponentiated within the C++ model, this
ensures that the estimated values of these parameters are always
positive.

```
# Set parameters to estimate
par <- list(
  logB0 = log(0.1), # Base rate of mass loss 
  logBf = log(0.1), # Mass loss increase in flight
  bd = 0, # Mean mass gain while diving
  b1 = 0, #  coefficient for effect of cov1 on G
  b2 = 0, # coefficient for effect of cov2 on G
  logSDP = log(sdp), # Process error - this is going to be a fixed parameter
  logA = log(0.25), # Constant (>0)
  logSDA = log(0.1), #  Log of parameter for the random effect on a [constant relating mass to WBF] (>0)
  logSDO = log(0.1),  # Observation error (>0)
  ai = rep(0, nrow(bid)), # a for each individual
  m = dat$mdata # starting values for mass
)
```

This creates a named list of factors mapping parameters that should
be should be estimated or fixed. See ?MakeADFun for a more detailed
explanation of parameter mapping.

As mentioned above, the map argument tells TMB the parameters are
estimated and those that are set to a fix value. The map values needs to
be factors, usually unique factor values. By default, TMB will assign
unique factor values to parameters. Importantly, if the map has a value
of NA for a parameter, TMB will use the value imputed as its starting
value as the value for that parameter; in other words TMB will not
estimate the parameter if the map value associated with it is NA. Here,
we fix the value of the standard deviation of the process equation. As
explained above we also fi the starting and end values for the predicted
masses.

```
# Map fixed values and random values
map <- list(logSDP = as.factor(NA), # fix sdp so it is not estimated by TMB
            m = as.factor(dat$mmap)#, # starting values for m, with NA for fixed start and end mass
            #ai = as.factor(1:nrow(bid))
)
```

We use TMB::MakeADFun() to construct the full model that combines
data, parameters, and mapped values with our c++ model. We use the
random argument to specify that m and ai are random effects.

```
# Set up model
mod <- MakeADFun(data = dataTmb, # list of all data objects required my the model
                 parameters = par, # list of all parameter objects required my the model
                 map = map, # list defining how to optionally collect and fix parameters
                 random = c("m", 'ai'), # defines random effect parameters
                 DLL = "wbf_mass_model", # name of loaded model object
                 inner.control = list(maxit=5000), # control values for inner optimization
                 tracepar = TRUE, 
                 silent=TRUE)
```

Run model and check convergence messages.

```
fit <- nlminb(mod$par, mod$fn, mod$gr, control = list(iter.max = 2000, eval.max=2000))

(fit$message)
```

```
## [1] "both X-convergence and relative convergence (5)"
```

##### Get estimates

Now we extract the parameter estimates from model fit using sd
report.

```
report <- summary(sdreport(mod)) # Extract estimated values from the model

param_est <- data.frame(report[c("b0","bf","bd", "b1", "b2","a","sda","sdo", "sdp"),]) 

param_est$ll <- param_est[,1] - (param_est[,2] * qnorm(0.05/2, lower.tail = F))
param_est$ul <- param_est[,1] + (param_est[,2] * qnorm(0.05/2, lower.tail = F))

param_est[c('bd','b1','b2'),c(1,3,4)] <- exp(param_est[c('bd','b1','b2'),c(1,3,4)])

round(param_est,3)
```

```
##     Estimate Std..Error    ll    ul
## b0     0.013      0.000 0.013 0.014
## bf     0.036      0.005 0.027 0.045
## bd     0.071      0.055 0.064 0.079
## b1     1.594      0.082 1.358 1.871
## b2     6.274      0.152 4.657 8.453
## a      0.249      0.001 0.246 0.252
## sda    0.011      0.001 0.009 0.013
## sdo    0.275      0.001 0.272 0.278
## sdp    0.100      0.000 0.100 0.100
```

Extract estimates of m (mass) and w (wbf). Calculate change in mass
(dm) and cumulative change in mass (dmass) through the deployment.

```
dat$m <- mod$report()$m
dat$w <- mod$report()$wpred
dat$w[dat$w < 1 | dat$w > 12] <- NA # take out non-flying observations

dat <- dat |> 
  dplyr::group_by(bird_id) |> 
  dplyr::mutate(
    dm = dplyr::lead(m) - m,
    dmass = cumsum(dm)
  )
```

##### Mass change within deployments

Extract data for an example trip.

```
coast <- rnaturalearth::ne_countries(scale = 10, returnclass = 'sf')

# Extract data from an example deployment

tt <- dat |> 
  dplyr::filter(bird_id == 6) |> 
  dplyr::mutate(
    dtime = as.numeric(difftime(time, min(time, na.rm = T), units = 'hours')),
    dmass = cumsum(dm)
  )

tt |> 
  dplyr::filter(trip_id == 1) |> 
  summarise(
    start_time = min(time),
    end_time = max(time),
    duration = round(as.numeric(difftime(end_time, start_time, units = 'hours')), 1),
    start_mass = round(m[1],1),
    end_mass = round(m[dplyr::n()],1),
    dmass = round(sum(dm),1),
    max_mass = max(m),
    peak_time = time[m == max_mass],
    end_forage = max(time[diving == 1]),
    return = min(time[flying == 1 & time > end_forage])
  )
```

```
## # A tibble: 1 × 11
##   bird_id start_time          end_time            duration start_mass end_mass
##     <int> <dttm>              <dttm>                 <dbl>      <dbl>    <dbl>
## 1       6 2023-07-07 19:26:20 2023-07-08 04:42:20      9.3      1027.    1095.
## # ℹ 5 more variables: dmass <dbl>, max_mass <dbl>, peak_time <dttm>,
## #   end_forage <dttm>, return <dttm>
```

The state-space model enables us to examine fine-scale changes in
mass within a deployment, to illustrate this we will focus on the mass
estimates from a single individual. The plots below depict changes in
mass spatially and temporally, as well as relative to distance from
colony and dive activity.

```
trip_sf <- sf::st_as_sf(tt, coords = c('lon','lat'), crs = 4326) # create simple features
track_sf <- trip_sf |> dplyr::summarise(do_union = FALSE) |> sf::st_cast('LINESTRING') # make linestring
bb <- c(-82.69078,  62.88172, -82.00538,  63.22442) # pretty bounding box

# map showing spatial distribution of mass changes within foraging trip
p1 <- ggplot() +
  geom_sf(data = coast) +
  geom_sf(data = trip_sf, aes(col = dm, size = dm), shape = 1) +
  scale_color_viridis_c(option = 'B') +
  scale_size_continuous(range = c(0.1, 3), guide = 'none') +
  coord_sf(xlim = bb[c(1,3)], ylim = bb[c(2,4)]) +
  scale_x_continuous(breaks = seq(-180, 180, 0.2)) +
  scale_y_continuous(breaks = seq(-180, 180, 0.1)) +
  labs(x ='', y = '', col = '\U0394 Mass (g)') +
  theme(
    text = element_text(size = 9),
    axis.text.y = element_text(angle = 90, hjust = 0.5),
    legend.position.inside = c(0.99, 0.99),
    legend.justification = c(1, 1),
    legend.background = element_blank(),
    legend.box = element_blank(),
    legend.key.size = unit(0.4, 'cm'),
  )
print(p1)
```

```
# ggsave('plots/Example trip map.png', plot = p1, units = 'in', width = 3.25, height = 3.25)
```

```
# plot showing cumulative change in mass over time since deployment and by behaviour, vertical lines show foraging trip start and end
p2 <- ggplot(tt, aes(x = dtime, y = m, col = behaviour, size = dm)) +
  geom_point() +
  geom_vline(xintercept = range(tt$dtime[!is.na(tt$trip_id)], na.rm = T), linetype = 2, linewidth = 0.5) +
  scale_color_viridis_d(option = 'D') +
  scale_size_continuous(range = c(0.1, 3), guide = 'none') +
  labs(x = 'Time since start (hr)', 
       y = 'Mass (g)', 
       col = '') +
  theme(
    text = element_text(size = 9),
    axis.text.y = element_text(angle = 90, hjust = 0.5),
    legend.position.inside = c(0.01, 0.01),
    legend.justification = c(0, 0),
    legend.background = element_blank(),
    legend.box = element_blank(),
    legend.key = element_blank(),
    legend.key.spacing = unit(0.01, 'cm'),
    legend.key.size = unit(0.4, 'cm')
  )

# plot showing change in mass with distance from colony, vertical lines show foraging trip start and end
p3 <- ggplot(tt, aes(x = dtime, y = coldist, col = dm, size = dm)) +
  geom_point() +
  geom_vline(xintercept = range(tt$dtime[!is.na(tt$trip_id)], na.rm = T), linetype = 2, linewidth = 0.5) +
  scale_color_viridis_c(option = 'B') +
  scale_size_continuous(range = c(0.1, 3), guide = 'none') +
  labs(x = 'Time since start (hr)', y = 'Distance from colony (km)',col = '\U0394 Mass (g)') +
  theme(
    text = element_text(size = 9),
    axis.text.y = element_text(angle = 90, hjust = 0.5),
    legend.position.inside = c(0.01, 0.99),
    legend.justification = c(0, 1),
    legend.background = element_blank(),
    legend.box = element_blank(),
    legend.box.background = element_blank(),
    legend.key = element_blank(),
    legend.key.size = unit(0.4, 'cm')
  )

# plot showing change in mass with dive activity, vertical lines show foraging trip start and end
p4 <- ggplot(tt) +
  geom_line(aes(x = dtime, y = depth_m), linewidth = 0.3) +
  geom_point(aes(x = dtime, y = depth_m, col = dm, size = dm)) +
  geom_vline(xintercept = range(tt$dtime[!is.na(tt$trip_id)], na.rm = T), linetype = 2, linewidth = 0.5) +
  scale_y_reverse() +
  scale_color_viridis_c(option = 'B') +
  scale_size_continuous(range = c(0.05, 3), guide = 'none') +
  labs(x = 'Time since start (hr)', y = 'Depth (m)',col = '\U0394 Mass (g)') +
  theme(
    text = element_text(size = 9),
    axis.text.y = element_text(angle = 90, hjust = 0.5, size = 8),
    legend.position.inside = c(0.01, 0.01),
    legend.justification = c(0, 0),
    legend.background = element_blank(),
    legend.box = element_blank(),
    legend.box.background = element_blank(),
    legend.key = element_blank(),
    legend.key.size = unit(0.4, 'cm')
  )
cowplot::plot_grid(p2, p3, p4, labels = c('A','B','C'), nrow = 3)
```

```
# ggsave('plots/Detailed foraging example.png', units = 'in', width = 6.5, height = 2.5*3)
```

##### Summarize trips

Estimating mass through the entire deployment allows us to calculate
the underlying mass gain for specific trips, even when a bird undertook
multiple trips or the initial deployment and recapture times do not
correspond well with trip start and end times. Here we summarize the
gross mass gain (i.e. mass change while foraging) and the net mass gain
(i.e. total change in mass) within each foraging trip in the case
study.

Foraging trips were defined as time spent away from the colony that
included:

- at least 15 min of diving
- lasted longer than 4 hours
- shorter than 24 hours

During incubation, adult murres usually alternate 12 hour shifts of
foraging with 12 hours at the colony. Trip duration constraints were
used to ensure we are only comparing regular foraging trips.

```
trips <- dat |> 
  dplyr::filter(!is.na(trip_id)) |> 
  dplyr::group_by(bird_id, trip_id) |> 
  dplyr::summarize(
    start = min(time),
    end = max(time),
    dur = dplyr::n()/(6 * 60),
    dive_time = sum(diving)/(6 * 60),
    max_depth = max(depth_m),
    fly_time = sum(flying)/(6 * 60),
    swim_time = sum(behaviour == 'swimming')/(6 * 60),
    coldist = max(coldist),
    start_mass = m[1],
    end_mass = m[dplyr::n()],
    max_mass = max(m),
    range_mass = max(m) - min(m),
    gross_gain = sum(dm[diving == 1]),
    net_gain = sum(dm),
    .groups = 'drop'
  ) |> 
  dplyr::filter(dive_time > 15/60, dur < 24, dur > 4) # filter on foraging trips
```

We can explore the distribution of gross mass gain and net mass gain
for all trips in the data. Because incubating murres alternate 12 hour
incubation bouts with 12 hour foraging bouts, we can calculate the
amount of mass a murre would need to cover losses while at the nest. By
comparing the net mass gain of foraging trips to the mass needed to
offset incubation costs, we can obtain a rough estimate of what
proportion of trips are successful enough to maintain long-term mass
balance.

```
# calculate break even point assuming a 12 hour incubation bout
bep <- 6*60*12 * report['b0',1]

# summarize gain during trips
gain_sum <- trips |> 
  dplyr::select(bird_id, trip_id, gross_gain, net_gain) |> 
  tidyr::pivot_longer(cols = c('gross_gain', 'net_gain'), names_to = 'measure', values_to = 'mass') |> 
  dplyr::group_by(measure) |> 
  dplyr::summarize(
    mean = mean(mass),
    sd = sd(mass),
    min = min(mass),
    max = max(mass),
    break_even = sum(mass > bep)/dplyr::n()
  )
gain_sum
```

```
## # A tibble: 2 × 6
##   measure     mean    sd   min   max break_even
##   <chr>      <dbl> <dbl> <dbl> <dbl>      <dbl>
## 1 gross_gain  81.8  34.5  12.8 160.       0.75 
## 2 net_gain    32.0  31.9 -40.8  99.3      0.208
```

```
grossgain_hist <- ggplot(trips) +
  geom_histogram(aes(gross_gain), bins = 10, fill = grey(0.9), col = 'black') +
  labs(x = 'Gross gain (g)', y = 'Number of trips') +
  theme(
    text = element_text(size = 9),
    axis.text.y = element_text(angle = 90, hjust = 0.5),
  )

netgain_hist <- ggplot(trips) +
  geom_histogram(aes(net_gain), bins = 10, fill = grey(0.9), col = 'black') +
  geom_vline(xintercept = mean(bep), linetype = 2) +
  labs(x = 'Net gain (g)', y = 'Number of trips') +
  theme(
    text = element_text(size = 9),
    axis.text.y = element_text(angle = 90, hjust = 0.5),
  )

cowplot::plot_grid(grossgain_hist, netgain_hist, labels = c('A','B'))
```

```
# ggsave('plots/Trip gain histograms.png', units = 'in', width = 6.5, height = 3.25)
```

Most trips result in a positive net mass gain. However, the average
mass lost during a 12 hour bout at the nest (57.1 g), only 75% of trips
exceed this threshold.

We now examine the relationships between net gain within a trip and
trip duration, maximum distance from the colony, and starting mass.

```
# relationship between trip duration and net mass gain 
mod1 <- lm(net_gain ~ dur, trips)
summary(mod1)
```

```
## 
## Call:
## lm(formula = net_gain ~ dur, data = trips)
## 
## Residuals:
##     Min      1Q  Median      3Q     Max 
## -71.435 -16.913   2.539  20.047  67.501 
## 
## Coefficients:
##             Estimate Std. Error t value Pr(>|t|)  
## (Intercept)  37.8544    17.7274   2.135   0.0381 *
## dur          -0.5738     1.6828  -0.341   0.7347  
## ---
## Signif. codes:  0 '***' 0.001 '**' 0.01 '*' 0.05 '.' 0.1 ' ' 1
## 
## Residual standard error: 32.17 on 46 degrees of freedom
## Multiple R-squared:  0.002521,   Adjusted R-squared:  -0.01916 
## F-statistic: 0.1163 on 1 and 46 DF,  p-value: 0.7347
```

```
nd <- data.frame(dur = seq(min(trips$dur), max(trips$dur), length.out = 100))
nd <- cbind(nd, predict(mod1, nd, interval = 'confidence'))

# plot the relationship
netgain_duration <- ggplot(trips) +
  geom_ribbon(data = nd, aes(x = dur, ymin = lwr, ymax = upr), fill = grey(0.9)) +
  geom_line(data = nd, aes(x = dur, y = fit)) +
  geom_point(aes(x = dur, y = net_gain)) +
  labs(x = 'Trip duration (hr)', y = 'Net gain (g)') +
  theme(
    text = element_text(size = 8),
    axis.text.y = element_text(angle = 90, hjust = 0.5),
  )
netgain_duration
```

The relationship with trip duration appears unimportant, and is not
statistically significant.

```
# relationship between maximum distance from the colony and net mass gain 
mod2 <- lm(net_gain ~ coldist, trips)
summary(mod2)
```

```
## 
## Call:
## lm(formula = net_gain ~ coldist, data = trips)
## 
## Residuals:
##     Min      1Q  Median      3Q     Max 
## -68.453 -18.243   1.457  19.838  73.432 
## 
## Coefficients:
##             Estimate Std. Error t value Pr(>|t|)  
## (Intercept)  12.3532     9.8325   1.256   0.2153  
## coldist       0.5646     0.2522   2.239   0.0301 *
## ---
## Signif. codes:  0 '***' 0.001 '**' 0.01 '*' 0.05 '.' 0.1 ' ' 1
## 
## Residual standard error: 30.59 on 46 degrees of freedom
## Multiple R-squared:  0.09824,    Adjusted R-squared:  0.07864 
## F-statistic: 5.011 on 1 and 46 DF,  p-value: 0.03006
```

```
nd <- data.frame(coldist = seq(min(trips$coldist), max(trips$coldist), length.out = 100))
nd <- cbind(nd, predict(mod2, nd, interval = 'confidence'))

# plot the relationship
netgain_dist <- ggplot(trips) +
  geom_ribbon(data = nd, aes(x = coldist, ymin = lwr, ymax = upr), fill = grey(0.9)) +
  geom_line(data = nd, aes(x = coldist, y = fit)) +
  geom_point(aes(x = coldist, y = net_gain)) +
  labs(x = 'Maximum distance from colony (km)', y = 'Net gain (g)') +
  theme(
    text = element_text(size = 8),
    axis.text.y = element_text(angle = 90, hjust = 0.5),
  )

netgain_dist
```

There appears to be a positive relationship between net gain and maximum
distance from colony of the trip.

```
# relationship between mass at the start of the trip and net mass gain 
mod3 <- lm(net_gain ~ start_mass, trips)
summary(mod3)
```

```
## 
## Call:
## lm(formula = net_gain ~ start_mass, data = trips)
## 
## Residuals:
##     Min      1Q  Median      3Q     Max 
## -74.061 -22.085   3.464  21.771  60.440 
## 
## Coefficients:
##              Estimate Std. Error t value Pr(>|t|)  
## (Intercept) 180.17065   77.89487   2.313   0.0252 *
## start_mass   -0.14771    0.07754  -1.905   0.0630 .
## ---
## Signif. codes:  0 '***' 0.001 '**' 0.01 '*' 0.05 '.' 0.1 ' ' 1
## 
## Residual standard error: 31.01 on 46 degrees of freedom
## Multiple R-squared:  0.07313,    Adjusted R-squared:  0.05298 
## F-statistic: 3.629 on 1 and 46 DF,  p-value: 0.06303
```

```
nd <- data.frame(start_mass = seq(min(trips$start_mass), max(trips$start_mass), length.out = 100))
nd <- cbind(nd, predict(mod3, nd, interval = 'confidence'))

# plot the relationship
netgain_start <- ggplot(trips) +
  geom_ribbon(data = nd, aes(x = start_mass, ymin = lwr, ymax = upr), fill = grey(0.9)) +
  geom_line(data = nd, aes(x = start_mass, y = fit)) +
  geom_point(aes(x = start_mass, y = net_gain)) +
  labs(x = 'Starting mass (g)', y = 'Net gain (g)') +
  theme(
    text = element_text(size = 8),
    axis.text.y = element_text(angle = 90, hjust = 0.5),
  )
netgain_start
```

There is a negative relationship between net gain and starting mass,
but it’s not statistically significant.

```
cowplot::plot_grid(netgain_duration, netgain_dist, netgain_start, labels = c('A','B',"C"), nrow = 1)
```

```
# ggsave('plots/Net gain regressions.png', units = 'in', width = 2.15 * 3, height = 2.15)
```

##### Map gain rate

To identify areas with the highest average mass gain while diving, we
summarized data to 1 hour intervals. Within each hour for each
deployment we calculated the gain rate while diving. This gain rate is
the sum of mass change while diving, divided by the time spent diving.
Only locations with at least 5 minutes of diving were included in the
analysis.

```
# Summarize mass gained at 1-hr intervals, based on median dive location
gr <- dat |> 
  dplyr::group_by(bird_id) |> 
  dplyr::mutate(
    time = lubridate::round_date(time, unit = '60 min') # summarize to 1 hour
  ) |> 
  dplyr::group_by(bird_id, time) |> 
  dplyr::summarise(
    gain_rate_ghr = sum(dm[behaviour == 'diving'])/(sum(diving) * 10/(60 * 60)), # gain rate while diving
    lon = median(lon[diving == 1]), # median longitude while diving
    lat = median(lat[diving == 1]), # median latitude while diving
    dive_time = ((sum(diving) * 10)/(60 * 60)), # total dive time in interval
    .groups = 'drop'
  ) |> 
  dplyr::filter(dive_time > 5/60) # filter locations with less than 5 min of diving
```

Convert locations to an equal area projection and extract coordinates
for spatial smoothing.

```
my_proj <- '+proj=laea +lon_0=-82.5206944 +lat_0=63.1010933 +datum=WGS84 +units=m +no_defs'
gr_sf <- sf::st_as_sf(gr, coords = c('lon','lat'), crs = 4326)
gr_sf <- sf::st_transform(gr_sf, crs = my_proj)
gr$x <- sf::st_coordinates(gr_sf)[,1]
gr$y <- sf::st_coordinates(gr_sf)[,2]

mapview::mapview(gr_sf, zcol = 'gain_rate_ghr')
```

To visualize spatial trends in mass gain, we performed a spatial
smooth on median hourly dive locations using the gain rate while diving
as the observed values. The spatial smooth was calculated on a 1x1 km
grid with 10 km buffer around observed locations. We used a distance of
5 km as the smoothing parameter.

Mass gain while diving generally increased from south to north. The
regions of highest gain were located north of the colony and west of
Bencas Island (not shown on this particular base map).

```
# buffer around observed locations (m)
b <- 10000

# mask areas with no data
r <- terra::rast(crs = my_proj, res = 1000, 
                 extent = c(c(min(gr$x) - b, max(gr$x) + b, min(gr$y) - b, max(gr$y) + b)))
m <- terra::rasterize(sf::st_buffer(gr_sf, b), r)

# create observation window for ppp
my_win <- owin(mask = data.frame(terra::crds(m)))

# create point pattern dataset
my_ppp <- ppp(x = gr$x,
              y = gr$y, 
              window=my_win,
              marks = gr$gain_rate_ghr)

# perform spatial smoothing 
my_sk <- Smooth(my_ppp, eps=1000, adjust = 1, edge = T, se = T, sigma = 5000)

# convert estimates to raster
r <- terra::rast(my_sk$estimate)
terra::crs(r) <- my_proj

# convert SE to raster
s <- terra::rast(my_sk$SE)
terra::crs(s) <- my_proj

# convert back to latlon for plotting
r <- terra::project(r, y = "epsg:4326")
s <- terra::project(s, y = "epsg:4326")

# extract values for ggplot
d <- data.frame(
  terra::crds(r, na.rm = T),
  values = terra::values(r, na.rm = T)[,1]
) 

# map raw observations
gr_proj <- sf::st_transform(gr_sf, crs = 4326)
gr1 <- ggplot() +
  geom_sf(data = gr_proj, aes(col = gain_rate_ghr), size = 2, alpha = 0.7) +
  geom_sf(data = coast, fill = grey(0.9)) +
  scale_colour_viridis_c(option = 'F', begin = 0.1, end = 0.8, direction = -1) +
  geom_point(aes(x = -82.01, y = 62.955), size = 3, col = 'blue') +
  coord_sf(xlim = terra::ext(r)[1:2], ylim = terra::ext(r)[3:4])+
  guides(color = guide_legend(reverse=FALSE)) +
  labs(x='', y='', colour = 'Gain rate (g/hr)') +
  theme(
    text = element_text(size = 8),
    axis.text.y = element_text(angle = 90, hjust = 0.5),
    legend.background = element_blank(),
    legend.box = element_blank(),
    legend.box.background = element_blank(),
    legend.key = element_blank(),
    legend.key.size = unit(0.4, 'cm')
  )

# map spatial smoothing estimates
bw <- 3
gr2 <- ggplot() +
  geom_contour_filled(data = d, aes(x = x, y = y, z = values), binwidth = bw) +
  geom_sf(data = coast, fill = grey(0.9)) +
  scale_fill_viridis_d(option = 'F', begin = 0.1, end = 0.8, direction = -1,
                       labels = paste0(seq(33, 60, bw), '-', seq(36, 63, bw))
  ) +
  geom_point(aes(x = -82.01, y = 62.955), size = 3, col = 'blue') +
  coord_sf(xlim = terra::ext(r)[1:2], ylim = terra::ext(r)[3:4]) +
  labs(x='', y='', fill = 'Gain rate (g/hr)') +
  theme(
    text = element_text(size = 8),
    axis.text.y = element_text(angle = 90, hjust = 0.5),
    legend.background = element_blank(),
    legend.box = element_blank(),
    legend.box.background = element_blank(),
    legend.key = element_blank(),
    legend.key.size = unit(0.4, 'cm')
  )
# ggsave('plots/Gain rate Est map.png', units = 'in', width = 3.25, height = 2.0)

# map standard error of smoothed estimates
gs <- data.frame(
  terra::crds(s, na.rm = F),
  values = terra::values(s)[,1]
) |> na.omit()

bw <- 0.6
gr3 <- ggplot() +
  geom_contour_filled(data = gs, aes(x = x, y = y, z = values), binwidth = bw) +
  geom_sf(data = coast, fill = grey(0.9)) +
  scale_fill_viridis_d(option = 'F', begin = 0.1, end = 0.8, direction = -1,
                       labels = paste0(seq(0, 3.6, bw), '-', seq(0.6, 4.2, bw))
  ) +
  geom_point(aes(x = -82.01, y = 62.955), size = 3, col = 'blue') +
  coord_sf(xlim = terra::ext(r)[1:2], ylim = terra::ext(r)[3:4]) +
  labs(x='', y='', fill = 'Uncertainty (SE)') +
  theme(
    text = element_text(size = 8),
    axis.text.y = element_text(angle = 90, hjust = 0.5),
    legend.background = element_blank(),
    legend.box = element_blank(),
    legend.box.background = element_blank(),
    legend.key = element_blank(),
    legend.key.size = unit(0.4, 'cm')
  )

cowplot::plot_grid(gr1, gr2, gr3, labels = c('A','B',"C"), ncol = 1)
```

```
# ggsave('plots/Gain rate map.png', units = 'in', width = 3.25, height = 2.0 * 3)
# ggsave('plots/Gain rate points - presentation.png', gr1, units = 'in', width = 5, height = 4)
# ggsave('plots/Gain rate smooth - presentation.png', gr2, units = 'in', width = 5, height = 4)
```

##### Compare model with no final mass

For the case study we fixed the final mass to measurements made on
the birds, which constrains the model to end at the known mass. We were
interested in exploring how much the parameter estimates and final mass
estimate would change if the final mass was unknown. Here we refit a
model without final mass.

Set up the data as if final mass is unknown. Create a new version of
`mdata` that does not include the final mass in the starting
values and remove NA from `mmap`.

```
# Starting values for mass estimates without including final mass
dat$mdata_nfm <- NA
dat$mdata_nfm[bid[,2] + 1] <- rdm0
dat$mdata_nfm <- imputeTS::na_locf(dat$mdata) # fill down with starting mass

# map input values for mass that do not fix final mass
dat$mmap_nfm <- 1:nrow(dat) # make sure all mmap values to be estimate are unique
dat$mmap_nfm[bid[,2] + 1] <- NA # fix start values to known starting mass

# Set parameters to estimate
par_nfm <- list(
  logB0 = log(0.1), # Base rate of mass loss 
  logBf = log(0.1), # Mass loss increase in flight
  bd = 0, # Mean mass gain while diving
  b1 = 0, #  coefficient for effect of cov1 on G
  b2 = 0, # coefficient for effect of cov2 on G
  logSDP = log(sdp), # Process error - this is going to be a fixed parameter
  logA = log(0.25), # Constant (>0)
  logSDA = log(0.1), #  Log of parameter for the random effect on a [constant relating mass to WBF] (>0)
  logSDO = log(0.1),  # Observation error (>0)
  ai=rep(0, nrow(bid)), # a for each individual
  m = dat$mdata_nfm # starting values for mass
)

# Map fixed values and random values
map_nfm <- list(logSDP = as.factor(NA), # fix sdp so it is not estimated by TMB
            m = as.factor(dat$mmap_nfm) # starting values for m, with NA for fixed start and end mass
)
```

Run new model without final mass.

```
# Set up model
mod_nfm <- MakeADFun(data=dataTmb, # list of all data objects required my the model
                 parameters=par_nfm, # list of all parameter objects required my the model
                 map = map_nfm, # ist defining how to optionally collect and fix parameters
                 random = c("m", 'ai'), # defines random effect parameters
                 DLL = "wbf_mass_model", # name of loaded model object
                 inner.control = list(maxit=5000), # control values for inner optimization
                 tracepar = TRUE, 
                 silent=TRUE)

fit_nfm <- nlminb(mod_nfm$par, mod_nfm$fn, mod_nfm$gr, control = list(iter.max = 2000, eval.max=2000))

(fit_nfm$message)
```

```
## [1] "relative convergence (4)"
```

Compare parameter estimates between the model fit with final mass and
without final mass.

```
report_nfm <- summary(sdreport(mod_nfm)) # Extract estimated values from the model

comp_est <- round(cbind(
  report[c("b0","bf","bd", "b1", "b2","a","sda","sdo", "sdp"),],
  report_nfm[c("b0","bf","bd", "b1", "b2","a","sda","sdo", "sdp"),]),
  4)

comp_est
```

```
##     Estimate Std. Error Estimate Std. Error
## b0    0.0132     0.0003   0.0156     0.0008
## bf    0.0360     0.0046   0.0384     0.0066
## bd   -2.6462     0.0554  -2.3879     0.0586
## b1    0.4662     0.0817   0.5866     0.0903
## b2    1.8364     0.1521   1.3426     0.1639
## a     0.2487     0.0015   0.2492     0.0015
## sda   0.0110     0.0011   0.0112     0.0011
## sdo   0.2752     0.0014   0.2725     0.0014
## sdp   0.1000     0.0000   0.1000     0.0000
```

There are some differences, but estimates are of the same magnitude
and direction.

This section calculates the RMSE between observed final mass and the
estimate from the model without fixed final mass.

```
dat$m_nfm <- mod_nfm$report()$m
dat$w_nfm <- mod_nfm$report()$wpred
dat$w_nfm[dat$w < 1 | dat$w > 12] <- NA # take out non-flying observations

dat <- dat |> 
  dplyr::group_by(bird_id) |> 
  dplyr::mutate(
    dm_nfm = dplyr::lead(m_nfm) - m_nfm,
    dmass_nfm = cumsum(dm_nfm)
  )

temp <- dat |> 
  dplyr::group_by(bird_id) |> 
  dplyr::summarize(
    start_mass = mass_on[1],
    end_mass = mass_off[1],
    end_nfm = m_nfm[dplyr::n()],
  )

ggplot(temp, aes(x = end_mass, y = end_nfm)) + 
  geom_point() +
  geom_abline(intercept = 0, slope = 1, linetype = 2) +
  labs(x = 'Observed final mass(g)', y = 'Estimated final mass(g)')
```

```
nfm_rmse <- Metrics::rmse(temp$end_mass, temp$end_nfm)
```

The RMSE for the difference between observed mass and mass estimated
from the model without final mass was 33.27. Plot below shows examples
of the full track differences in mass estimates from the model fit with
final mass (black) and the model fit without final mass (red).

```
dat |> 
  dplyr::filter(bird_id %in% 0:11) |> 
  ggplot() +
  geom_line(aes(x = time, y = m), col = 'black') +
  geom_line(aes(x = time, y = m_nfm), col = 'red') +
  ylim(range(c(dat$m, dat$m_nfm))) +
  labs(x = 'Time', y = 'Mass (g)') +
  scale_x_datetime(date_labels = '%H:%M') +
  facet_wrap(.~bird_id, nrow = 3, scales = 'free') +
  theme(
    text = element_text(size = 8)
  )
```
