## Supplementary material for "Using wingbeat frequency to estimate mass gained by seabirds": simulations.Rmd

````
---
title: "Using wing beat frequency to estimate mass gained by seabirds. <br> Supplementary material C: simulation analyses"
date: "`r Sys.Date()`"
output: html_document
---

```{r setup, include=FALSE}
knitr::opts_chunk$set(echo = TRUE)
```


# General set-up

## Load packages and basic plot set up

```{r, message=FALSE}
library(TMB)
library(tidyr)
library(dplyr)
library(ggplot2)
library(ggpubr)
library(latex2exp) # For math symbol in plots
theme_set(theme_light())
```

## Compile C++ Models

Compile and load the c++ models.

The file **`wbf_mass_model.cpp`** contains the function to calculate the negative log likelihood of the case study model presented in the main text.

```{r compile, message=FALSE}
compile("wbf_mass_model.cpp") # Compile c++ code
dyn.load(dynlib("wbf_mass_model")) # Load model in R
```

# Setting up all functions needed for the simulation analyses

Here, we are setting up all the functions that we need to run the simulation analyses. We also perform a quick test that these functions return the results we expect.

## Function to simulate data from behavioural states

Function to simulate data using the same model as the one used in the case study in the main text.

```{r sim-fun, cache=TRUE}
sim_data <- function(data, m0, par) {
  # data: list, with each element containing a data frame with the data for one individual. 
  #       Time series of flying, diving, wbf, cov1, and cov2.
  # m0: list of started mass value for each individual
  # par: parameters of the model
  alpha = par[1] # Constant for relating wbf to mass - alpha
  sigma_m = par[2] # Error term for process equation - sigma_m
  sigma_w = par[3] # Error term for observation equation (WBF) - sigma_w
  beta_0 = par[4] # Base rate for mass loss over time - beta_0
  beta_f = par[5] # Increased mass loss during flight - beta_f
  beta_d = par[6] # Base mass gain while diving - beta_d
  sigma_a = par[7] # Individual variation in a - sigma_a
  beta_1 = par[8] # Effect of covariate 1 on mass gain while diving - beta_1
  beta_2 = par[9] # Effect of covariate 2 on mass gain while diving - beta_2
  
  ai <- rep(NA, length(data))
  
  # loop for each individual
  for (j in 1:length(data)) {
    
    ###### Process equation
    # Empty vector to put mass values
    mass <- rep(NA, nrow(data[[j]])+1)
    # Starting mass value is known
    mass[1] <- m0[j]
    
    # For all other time steps, use process equation
    for (i in 2:length(mass)) {
      mass[i] <- 
        mass[i-1] - beta_0  - # previous mass - baseline loss
        (data[[j]]$flying[i - 1] * beta_f) + # extra loss if flying
        # gain if diving, a function of 2 covariates
        (data[[j]]$diving[i-1]*exp(beta_d + beta_1*data[[j]]$cov1[i-1] + 
                                  beta_2*data[[j]]$cov2[i-1])) + 
        rnorm(1, 0, sigma_m)
    }
    
    ###### Observation equation
    
    # Random effect for a
    ai[j] <- rnorm(1, alpha, sigma_a)
    
    
    for (i in 1:(length(mass)-1)){
      # If flying, link mass to wing beat frequency
      if (data[[j]]$flying[i] == 1){ 
        data[[j]]$wbf[i] <- ai[j] * sqrt(mass[i+1]) + rnorm(1, 0, sigma_w)
      } 
    }
    
    data[[j]]$mass <- mass[-1]
    data[[j]]$ai <- ai[j]
    if (min(data[[j]]$mass, na.rm = T) < 0) {
      stop("minimum mass less than 0! try changing sim values")
    }
  }
  
  return(data)
}
```

## Function to read in data used to simulate mass and wing beat frequency

Here, we make a function to extract the real bird data that we use for simulation. Specifically, we use time flying, diving, and mass at start and end. We also extract the covariates from the real bird data. Specifically, max depth of a dive and the standard deviation of pitch, both of which are strictly positive. As in the case study, both are scaled by dividing their values by the maximum value observed for each.

``` {r murre-data-fx, cache=TRUE}
extract_murre_data_4_sim <- function(file_name = "case_study_data.csv", sub_birds = NULL){

    all_data <- read.csv(file_name)
    
    if(is.null(sub_birds)){
      sub_birds <- 1:length(unique(all_data$bird_id))
    }
    
    rd <- all_data[all_data$bird_id %in% unique(all_data$bird_id)[sub_birds],]
    
    
    ### Covariates
    # Same covariates that are used in the case study

    rd$cov1 <- rd$max_depth_m/max(rd$max_depth_m)
    rd$cov2 <- rd$pitch_sd/max(rd$pitch_sd)

    # Starting mass for all individuals as a list
    rdm0 <- unlist(lapply(unique(rd$bird_id), 
                      FUN = function(x) head(rd$mass_on[rd$bird_id == x], 1)))

    # Final mass for all individuals as a list
    rdmT <- unlist(lapply(unique(rd$bird_id), 
                      FUN = function(x) tail(rd$mass_off[rd$bird_id == x], 1)))
    
    # Convert to list of data frames
    dd <- unique(rd$bird_id)
    rd <- lapply(unique(rd$bird_id), 
             FUN = function(x) rd[rd$bird_id == x,
                                  c("flying", "diving", "wbf","cov1","cov2")])
    names(rd) <- dd

  
   return(list(all_data = all_data, rd = rd, rdm0 = rdm0, rdmT = rdmT,
               sub_birds = sub_birds)) 
} 
```


## Function to prepare the data for TMB

Function to transform the data from simulation in a format for TMB functions.

``` {r prep-TMB-fx, cache=TRUE}
prep_data <- function(sim_data_raw, all_data, sub_birds, rdm0, rdmT){
  
  temp <- do.call(rbind, sim_data_raw)
  rd <- all_data[all_data$bird_id %in%
                   unique(all_data$bird_id)[sub_birds],]
  temp$bird_id <- as.numeric(as.factor(rd$bird_id)) - 1
  temp$idx <- 0:(nrow(temp) - 1)
  
  bid <- temp |> 
    dplyr::group_by(bird_id) |> 
    dplyr::summarize(
      start = min(idx),
      end = max(idx)
  ) 
  
  rdm0 <- temp$mass[bid$start + 1]
  rdmT <- temp$mass[bid$end + 1]
  
  temp$zdata <- NA
  
  # Starting values for mass, to help TMB
  temp$zdata[(bid$start) + 1] <- rdm0
  temp$zdata[(bid$end) + 1] <- rdmT
  temp$zdata <- imputeTS::na_locf(temp$zdata)
  
  # fix start and end mass
  temp$zmap <- 1:nrow(temp) # Important do not make factor here
  temp$zmap[bid$start + 1] <- NA
  temp$zmap[bid$end + 1] <- NA
  
  # Important: if there are additional levels to the factor, you'll run into problems!
  # So make sure not to make levels before adding NAs!

  bid <- as.matrix(bid)
  
  # data object to pass to TMB
  dataTmb <- list(w = temp$wbf, # Skip first record of WBF
                d = temp$diving, # Include all diving
                f = temp$flying,
                bid = bid,
                c1 = temp$cov1,
                c2 = temp$cov2
                ) # Include all flying
  return(list(dataTmb = dataTmb, 
                    zinfo = temp[,c("zdata", "zmap", "mass")]))
}
```


## Function to fit the model

Function to fit the model. It calls in all of the data needed to do so.

``` {r setup-mod-fx, cache=TRUE}
fit_mod_tmb <- function(par0, tmb_data_z, sub_birds, end_mass_inc = TRUE, rel.tol = NULL){
  # Extract info from tmb_data_z
  dataTmb <- tmb_data_z$dataTmb
  zinfo <- tmb_data_z$zinfo
  
  # Set parameters to estimate
  par2pTmb <- list(logA = log(par0[1]), 
                 logSDP = log(par0[2]), 
                 logSDO = log(par0[3]), 
                 logB0 = log(par0[4]), 
                 logBf = log(par0[5]),
                 bd = par0[6],
                 logSDA = log(par0[7]),
                 b1 = par0[8],
                 b2 = par0[9],
                 m = zinfo$zdata,
                 ai = rep(0, length(sub_birds))
                 ) 
  
  
  if(!end_mass_inc){
    zinfo$zmap[length(zinfo$zmap)] <- zinfo$zmap[length(zinfo$zmap)-1] + 1 
  }
  
    # Map fixed values
    # One fix paramater: sigma_m
    mapTmb <- list(logSDP = as.factor(NA),
    # Fix both start and end mass values (NA in zinfo$zmap)
        m = as.factor(zinfo$zmap),
    # Not fixed, one a_i for each individual
        ai = as.factor(1:length(sub_birds)))

  # Set up TMB model
  mod <- MakeADFun(data = dataTmb, 
                    parameters = par2pTmb, 
                    random = "m",
                    map = mapTmb, 
                    DLL = "wbf_mass_model",
                    inner.control = list(maxit=5000),
                    tracepar = TRUE, 
                    silent=TRUE)
  
  mod$env$tracepar <- FALSE
  
  # Fit model
  if(is.null(rel.tol)){
    fit <- nlminb(mod$par, mod$fn, mod$gr, 
              control = list(iter.max = 2000, eval.max=2000))  
  }else{
    fit <- nlminb(mod$par, mod$fn, mod$gr, 
              control = list(iter.max = 2000, eval.max=2000, rel.tol=rel.tol))  
  }
  

  return(list(fit = fit, mod = mod))
}
```

## Test out functions on small simulation

Here, we will use a small dataset to test out the functions we have created above for the simulation analyses.
First, we will extract the data needed for the simulations. In our simple test here, we only look at the first 10 individuals in the murre dataset.

```{r}
murre_data <- extract_murre_data_4_sim(sub_birds = 1:10)
```

Simulate mass data using the parameter values fixed and estimated in the case study (see Table 1).

``` {r run-sim, cache=TRUE}
# Simulated parameter values
sim_par <- c(alpha = 0.249, sigma_m = 0.1, sigma_w = 0.275, beta_0 = 0.013, 
             beta_f = 0.036, beta_d = -2.646, sigma_a = 0.011, 
             beta_1 = 0.466, beta_2 = 1.836)


# Simulate data
set.seed(123)
temp <- sim_data(data = murre_data$rd, m0 = murre_data$rdm0, par = sim_par)
```

Plot the results to quickly check that all is good.

```{r plot_sim_test, cache=TRUE, warning=FALSE}
# Plot simulated data
for (i in 1:length(temp)) {
  p <- ggplot(temp[[i]]) + 
    geom_point(aes(x = 1:nrow(temp[[i]]), y = mass, col = diving)) +
    geom_point(aes(x = 1:nrow(temp[[i]]), y = ((wbf/ai[1])^2)), col = 'red') +
    labs(x = 'Time', y = 'Mass (g)')
  print(p)
}
```

Transform the simulated data from `temp` in a format that can be handed to TMB as vectors.

``` {r prep-TMB, cache=TRUE, message=FALSE}
tmb_data_z <- prep_data(temp, murre_data$all_data, murre_data$sub_birds, 
                        murre_data$rdm0, murre_data$rdmT)
``` 

Fit model on the simulated data.

I note that the model is so complex that we often run into convergence problems, and we need decrease the relative tolerence and therefore the precision of the estimates

**Note: this will take a few minutes.** 

```{r run-sim-data, cache=TRUE}
sim_fit_1 <- fit_mod_tmb(sim_par, 
                         tmb_data_z, murre_data$sub_birds, rel.tol = 1e-8)
sim_fit_1$fit$message
```

Some random seed will result in false convergence messages, and thus we will need to account for this in the full simulation analyses.

Here, we assess whether the parameter estimates are close to those used to simulate the data.

``` {r check-sim-est, cache=TRUE}
sdrmod <- summary(sdreport(sim_fit_1$mod))

cbind(sdrmod[c("a", "sdp", "sdo","b0","bf","bd",
               "sda","b1","b2"),], sim_par[1:9])
```

These look good, some parameters are a bit off, but the estimate are in the same order of magnitude as those used in the simulation. The full simulation studies below will properly assess whether there are biases.

# Simulation study 1: sensitivity of results to $\sigma_m$

State-space models can have difficulty simultaneously estimating the parameters associated with process stochasticity and measurement error (in our case $\sigma_m$ and $\sigma_w$; Auger-Méthé et al. 2016, 2021). 

We noticed with our case study that we had convergence issues and problems when we estimated both, and we decided as such to fix the standard deviation associated with the process stochasticity, $\sigma_m$, to 0.1, to limit this problem. We chose this value as we expect approximately 95% of the additional changes in murre mass in 10 seconds to fall with - 0.2 and 0.2 g.

While this is a biologically relevant value, the exact value is arbitrary and we wanted to do a sensitivity analysis to verify how robust our results are when $\sigma_m$ is misspecified.

To do so, we will simulate 10 individuals, using three different $\sigma_m$ values: $(0.05, 0.1, 0.15)$. We will change the $\sigma_m$ used in the simulations, and the $\sigma_m$ fixed when we fit the model, we will look at all combinations of simulated and fixed $\sigma_m$ values for a total of 9 combinations. For the other parameter values, we are using the values estimated in our case study: $(\beta_0 = 0.013, \beta_f = 0.036, \beta_d = -2.646, \beta_{depth} = 0.466, \beta_{pitch} = 1.836, \alpha = 0.249, \sigma_w = 0.275, \sigma_a = 0.011)$. We repeat each of these scenarios 100 times, and compute the root mean square error for the state and parameter values.

```{r par-sa, cache = TRUE}
n_sim <- 100
n_i <- 20
sigma_m_seq_sim <- c(0.05, 0.1, 0.15)
sigma_m_seq_fitted <- c(0.05, 0.1, 0.15)
```

**Note: this will take multiple hours/days to run on most computers.**

```{r sa-sim, cache=TRUE, warning=FALSE}
sa_res <- data.frame(matrix(
  nrow = n_sim*length(sigma_m_seq_sim)*length(sigma_m_seq_fitted), 
  ncol = 13))
colnames(sa_res) <- c("sigma_m_sim", "sigma_m_fitted", "sim_index", 
                      "alpha", "sigma_w", "beta_0", "beta_f", "beta_d", 
                      "sigma_a", "beta_1", "beta_2", "RMSE_m",
                      "message")
set.seed(1234) 
i <- 1 # index
t0 <- Sys.time()
for(s in 1:length(sigma_m_seq_sim)){
  for(j in 1:n_sim){
    # Simulated parameter values
    sim_par_sa_1 <- c(alpha = 0.249, sigma_m = sigma_m_seq_sim[s], sigma_w = 0.275, 
                    beta_0 = 0.013, beta_f = 0.037, beta_d = 0.058, sigma_a = 0.011, 
                    beta_1 = 0.070, beta_2 = 0.231)
    
    # Simulate data
    bid_sa_1 <- sort(sample(1:55, n_i))
    murre_data_sa_1 <- extract_murre_data_4_sim(sub_birds = bid_sa_1)
    sim_sa_1 <- sim_data(data = murre_data_sa_1$rd, 
                       m0 = murre_data_sa_1$rdm0, par = sim_par_sa_1)
  
    # Fit with tmb
    sim_data_z_sa_1 <- prep_data(sim_sa_1, murre_data_sa_1$all_data, 
                             murre_data_sa_1$sub_birds, 
                             murre_data_sa_1$rdm0, murre_data_sa_1$rdmT)

    fit_par0_sa_1 <- sim_par_sa_1
    for(k in 1:length(sigma_m_seq_fitted)){
      sa_res[i, "sigma_m_sim"] <- sigma_m_seq_sim[s]
      sa_res[i, "sim_index"] <- i
      sa_res[i, "sigma_m_fitted"] <- sigma_m_seq_fitted[k]
      
      fit_par0_sa_1["sigma_m"] <- sigma_m_seq_fitted[k]
      sim_fit_sa_1 <- fit_mod_tmb(fit_par0_sa_1, sim_data_z_sa_1, 
                            murre_data_sa_1$sub_birds, 
                            rel.tol = 1e-8)
      sa_res[i, "message"] <- sim_fit_sa_1$fit$message
      
      # Parameter estimates
      sim_par_est_sa_1 <- summary(sdreport(sim_fit_sa_1$mod))
      sa_res[i, 
           c("alpha", "sigma_w", "beta_0", "beta_f", "beta_d", 
                      "sigma_a", "beta_1", "beta_2")] <- 
        sim_par_est_sa_1[c("a", "sdo","b0","bf","bd",
               "sda","b1","b2"), "Estimate"]
      
      # RMSE - mass
      mass_pred_sa_1 <- sim_par_est_sa_1[row.names(sim_par_est_sa_1)=="m", "Estimate"]
      sim_mass_sa_1 <- sim_data_z_sa_1$zinfo$mass[!is.na(sim_data_z_sa_1$zinfo$zmap)]
      sa_res[i, "RMSE_m"] <- 
        sqrt(sum((sim_mass_sa_1 - mass_pred_sa_1)^2)/length(sim_mass_sa_1))
    
      i <- i + 1
    }
  }
}
Sys.time() - t0 # 16 min for 1 sim 20 individuals 
# Estimated time: (16*100)/60
```

Note that running this code will results in some warnings of the type `Warning in nlminb ... NA/NaN function evaluation`, that are not printed here to reduce the length of the document.

Save data

```{r sa_write, cache=TRUE}
write.csv(sa_res, file = paste("sa_res_", format(Sys.time(), "%Y%m%d"), ".csv",
                               sep=""), 
          row.names = FALSE)
```

```{r sa_read, eval=FALSE, echo = FALSE}
# Read data 
# sa_res <- read.csv("sa_res_20250202.csv") # 20 ind.
# sim_par_sa_1 <- c(alpha = 0.249, sigma_m = NA, sigma_w = 0.275,
#                   beta_0 = 0.013, beta_f = 0.037, beta_d = 0.058, sigma_a = 0.011,
#                   beta_1 = 0.070, beta_2 = 0.231)

```

Look at results

Check convergence

```{r sa_conv_mess, cache=TRUE}
sa_res %>%
  group_by(sigma_m_sim, sigma_m_fitted, message) %>%
  summarize(n_mess = n(), .groups = "keep")
```

```{r sa_conv, cache=TRUE}
unique(sa_res$message)

conv_prob <- sa_res %>% filter(message == "false convergence (8)" | message == "singular convergence (7)")
nrow(conv_prob)/nrow(sa_res)
```

A large proportion of models had convergence problems, which indicated that when analyzing a real dataset, we may want to pay close attention to convergence issues. Here, we explore the simulations that reached at least relative convergence.

```{r sa_conv_rel, cache=TRUE}
sa_res_converged <- sa_res %>% filter(message != "false convergence (8)" & message != "singular convergence (7)")
```

```{r sa_res, cache=TRUE}
# Colour blind friendly palette: IBM
col_sigma <- c("#648FFF", "#DC267F", "#FFB000")


RMSE_plot <- ggplot() + 
  geom_boxplot(data = sa_res_converged,
               aes(x = as.factor(sigma_m_sim), y = RMSE_m, 
                   col = as.factor(sigma_m_fitted),
                   fill = as.factor(sigma_m_fitted)),
               alpha = 0.6) +
  scale_color_manual(name = TeX(r"($\sigma_m$ of fitted model)"), 
                     values = col_sigma) +
  scale_fill_manual(name = TeX(r"($\sigma_m$ of fitted model)"), 
                     values = col_sigma) +
  xlab(TeX(r"($\sigma_m$ of simulations)")) +
  ylab("RMSE") + 
  theme(legend.position="top")
RMSE_plot
```

As expected, the size of the difference between the simulated and predicted values increases when we use the wrong values in the model. However, the magnitude of these additional errors is small compared to how much the error increases as a function of the underlying $\sigma_m$. In essence, if the mass of the animals is very variable (high true $\sigma_m$ values - here high simulation $\sigma_m$ values), it is harder to predict the true mass than when $\sigma_m$ is small, and the effect of misspecification is small in comparison. 

Plot of parameters

```{r par_plot_function, cache=TRUE}
par_plot <- function(res, par_name, TeX_par, col){
  column <- sym(par_name)
  p_plot <- ggplot() + 
    geom_boxplot(data = res,
                 aes(x = as.factor(sigma_m_sim), y = !!column,
                     col = as.factor(sigma_m_fitted),
                     fill = as.factor(sigma_m_fitted)),
                 alpha = 0.6,
                 outlier.size = 0.5,
                 lwd = 0.2) +
  scale_color_manual(name = TeX(r"($\sigma_m$ of fitted model)"), 
                     values = col_sigma) +
  scale_fill_manual(name = TeX(r"($\sigma_m$ of fitted model)"), 
                     values = col_sigma) +
    geom_hline(yintercept = sim_par_sa_1[par_name], col = col) +
    xlab(TeX(r"($\sigma_m$ of simulations)")) +
    ylab(TeX_par)
  return(p_plot)
}
```

Plot for each parameter.

```{r sa_par_plot, cache=TRUE}
sigma_w_plot <- par_plot(sa_res_converged, "sigma_w", TeX(r"($sigma_w$)"), grey(0.3))
alpha_plot <- par_plot(sa_res_converged, "alpha", TeX(r"($alpha$)"), grey(0.3))
sigma_a_plot <- par_plot(sa_res_converged, "sigma_a", TeX(r"($sigma_a$)"), grey(0.3))
beta_0_plot <- par_plot(sa_res_converged, "beta_0", TeX(r"($beta_0$)"), grey(0.3))
beta_f_plot <- par_plot(sa_res_converged, "beta_f", TeX(r"($beta_f$)"), grey(0.3))
beta_d_plot <- par_plot(sa_res_converged, "beta_d", TeX(r"($beta_d$)"), grey(0.3))
beta_1_plot <- par_plot(sa_res_converged, "beta_1", TeX(r"($beta_1$)"), grey(0.3))
beta_2_plot <- par_plot(sa_res_converged, "beta_2", TeX(r"($beta_2$)"), grey(0.3))
```


Plot the parameter plots together.

```{r sigma_sim_par, fig.height = 5, fig.width = 7}
ggarrange(sigma_w_plot, sigma_a_plot, alpha_plot, beta_0_plot,
          beta_f_plot, beta_d_plot, beta_1_plot, beta_2_plot,
          nrow = 2, ncol = 4, common.legend = TRUE,
          labels = LETTERS[1:8],
          font.label = list(size = 10),
          hjust = -1.2, vjust = 2,
          align ="hv") 
```

# Simulation study 2: importance of including final mass

In the case study, since we re-weighed the individual when we remove their tag, we fixed the value of the mass at the end of the time series. Doing so helped model fitting, and we wanted to demonstrate the effect of including the final mass by running simulations with and without that final mass included.

Here we use the a $\sigma_m$ value of 0.1 for the simulation and the value fixed in the model. For the parameters that are estimated, we use the estimated values from the case study for the simulations and as starting values.

```{r par-fix-end-mass, cache = TRUE}
n_sim <- 100
n_i <- 20

# Simulated/fix/par0 parameter values
sim_par_mass_f <- c(alpha = 0.249, sigma_m = 0.1, sigma_w = 0.275, 
                    beta_0 = 0.013, beta_f = 0.036, beta_d = -2.646, sigma_a = 0.011, 
                    beta_1 = 0.466, beta_2 = 1.836)
```

This is a similar set up as previous simulation analysis.

Note that we remove the final mass in RMSE when mass is not fixed to have comparable RMSE.

**Note: this will take multiple hours to run on most computers.**

```{r mass_fixed_sim, cache=TRUE, message=FALSE, warning=FALSE}
mass_fix_res <- data.frame(matrix(
  nrow = n_sim*2, 
  ncol = 12))
colnames(mass_fix_res) <- c("end_mass_fix", "sim_index", 
                      "alpha", "sigma_w", "beta_0", "beta_f", "beta_d", 
                      "sigma_a", "beta_1", "beta_2", "RMSE_m",
                      "message")
set.seed(1234) 
t0 <- Sys.time()
i <- 1
for(j in 1:n_sim){
    
    
    # Simulate data
    bid_mass_f <- sort(sample(1:55, n_i))
    murre_data_mass_f <- extract_murre_data_4_sim(sub_birds = bid_mass_f)
    sim_mass_f <- sim_data(data = murre_data_mass_f$rd, 
                       m0 = murre_data_mass_f$rdm0, par = sim_par_mass_f)
  
    # Fit with tmb
    sim_data_z_mass_f <- prep_data(sim_mass_f, murre_data_mass_f$all_data, 
                             murre_data_mass_f$sub_birds, 
                             murre_data_mass_f$rdm0, 
                             murre_data_mass_f$rdmT)
    
    

    # End mass fixed
    mass_fix_res[i, "sim_index"] <- j
    mass_fix_res[i, "end_mass_fix"] <- TRUE
    sim_fit_mass_f_y <- fit_mod_tmb(sim_par_mass_f, sim_data_z_mass_f, 
                            murre_data_mass_f$sub_birds,
                            end_mass_inc = TRUE, 
                            rel.tol = 1e-8)
    
    mass_fix_res[i, "message"] <- sim_fit_mass_f_y$fit$message
    
    # Parameter estimates
    sim_par_est_mass_f_y <- summary(sdreport(sim_fit_mass_f_y$mod))
    
    mass_fix_res[i, 
           c("alpha", "sigma_w", "beta_0", "beta_f", "beta_d", 
                      "sigma_a", "beta_1", "beta_2")] <- 
        sim_par_est_mass_f_y[c("a", "sdo","b0","bf","bd",
               "sda","b1","b2"), "Estimate"]
      
    
      
      # RMSE - mass
      mass_pred_mass_f_y <- sim_par_est_mass_f_y[row.names(sim_par_est_mass_f_y)=="m", "Estimate"]
      sim_mass_mass_f_y <- sim_data_z_mass_f$zinfo$mass[!is.na(sim_data_z_mass_f$zinfo$zmap)]
      mass_fix_res[i, "RMSE_m"] <- 
        sqrt(sum((sim_mass_mass_f_y - mass_pred_mass_f_y)^2)/length(mass_pred_mass_f_y))
    
      i <- i + 1
      
      
      # End mass not fixed
      mass_fix_res[i, "sim_index"] <- j
      mass_fix_res[i, "end_mass_fix"] <- FALSE
      sim_fit_mass_f_n <- fit_mod_tmb(sim_par_mass_f, sim_data_z_mass_f, 
                            murre_data_mass_f$sub_birds, 
                            end_mass_inc = FALSE, 
                            rel.tol = 1e-8)
      
      mass_fix_res[i, "message"] <- sim_fit_mass_f_n$fit$message
      
      # Parameter estimates
      sim_par_est_mass_f_n <- summary(sdreport(sim_fit_mass_f_n$mod))
      mass_fix_res[i, 
           c("alpha", "sigma_w", "beta_0", "beta_f", "beta_d", 
                      "sigma_a", "beta_1", "beta_2")] <- 
        sim_par_est_mass_f_n[c("a", "sdo","b0","bf","bd",
               "sda","b1","b2"), "Estimate"]
      
    
      
      # RMSE - mass
      mass_pred_mass_f_n <- sim_par_est_mass_f_n[row.names(sim_par_est_mass_f_n)=="m", "Estimate"]
      # Exclude final mass, to make comparison fair
      mass_pred_mass_f_n <- mass_pred_mass_f_n[-length(mass_pred_mass_f_n)]
      # length(mass_pred_mass_f_n)
      sim_mass_mass_f_n <- sim_data_z_mass_f$zinfo$mass[!is.na(sim_data_z_mass_f$zinfo$zmap)]
      # length(sim_mass_mass_f_n)
      mass_fix_res[i, "RMSE_m"] <- 
        sqrt(sum((sim_mass_mass_f_n - mass_pred_mass_f_n)^2)/length(mass_pred_mass_f_n))
      
      i <- i + 1
}
Sys.time() - t0 # 6.3 min for 1 sim 20 individuals 
# Estimated time: (6.3*100)/60
```

Note that running this code will results in some warnings of the type `Warning in nlminb ... NA/NaN function evaluation`, that are not printed here to reduce the length of the document.

Save data

```{r mass_fix_write, cache=TRUE}
write.csv(mass_fix_res, file = paste("mass_fix_res_", format(Sys.time(), "%Y%m%d"), ".csv",
                               sep=""), 
          row.names = FALSE)
```

```{r mass_read, eval=FALSE, echo = FALSE}
# Read data 
mass_fix_res <- read.csv("mass_fix_res_20250203.csv") # 20 ind.
```


Here, we look at the convergence messages.

```{r mass_f_conv_mess, cache=TRUE}
mass_fix_res %>%
  group_by(end_mass_fix,message) %>%
  summarize(n_mess = n(), .groups = "keep")
```

```{r}
mass_fix_res_conv <- mass_fix_res %>% 
  filter(message == "relative convergence (4)")

1-nrow(mass_fix_res_conv)/nrow(mass_fix_res)
```

```{r mass_f_rmse, cache=TRUE}
# Colour blind friendly
col_included <- c("#E1BE6A", "#40B0A6")
RMSE_plot_mass <- ggplot() + 
  geom_boxplot(data = mass_fix_res_conv,
               aes(x = end_mass_fix, y = RMSE_m,
                   col = end_mass_fix,
                   fill = end_mass_fix),
               alpha = 0.6, show.legend = FALSE) +
  scale_color_manual(name = "", 
                     values = col_included) +
  scale_fill_manual(name = "", 
                     values = col_included) +
  xlab("Final mass included?") +
  ylab("RMSE") +
  scale_x_discrete(labels=c("FALSE" = "No", "TRUE" = "Yes"))
RMSE_plot_mass
```


Plot function for parameters.

```{r mass-alpha-plot, cache=TRUE}
par_plot_mass <- function(res, par_name, TeX_par, col = grey(0.3)){
  column <- sym(par_name)
  # scale of y-axis
  yaxis <- res %>% select(!!column) %>% range()
  yaxis[1] <- floor(yaxis[1] * 1000)/1000
  yaxis[2] <- ceiling(yaxis[2] * 1000)/1000
  
  p_plot <- ggplot() + geom_boxplot(data = res,
               aes(x = end_mass_fix, y = !!column,
                     col = end_mass_fix,
                     fill = end_mass_fix),
                 alpha = 0.6,
                 outlier.size = 0.5,
                 lwd = 0.2, show.legend = FALSE) +
    scale_color_manual(name = "Included", 
                     values = col_included ) +
    scale_fill_manual(name = "Included", 
                     values = col_included) +
    geom_hline(yintercept = sim_par_mass_f[par_name], col = col) +
    xlab("Final mass inc.") +
    ylab(TeX_par) +
    ylim(yaxis) +
    scale_x_discrete(labels=c("FALSE" = "No", "TRUE" = "Yes"))
  
  return(p_plot)
}
```

Create each parameter plot.

```{r mass_f_par_plots, cache = TRUE}
sigma_w_plot_m <- par_plot_mass(mass_fix_res, "sigma_w", TeX(r"($sigma_w$)"))
alpha_plot_m <- par_plot_mass(mass_fix_res, "alpha", TeX(r"($alpha$)"))
sigma_a_plot_m <- par_plot_mass(mass_fix_res, "sigma_a", TeX(r"($sigma_a$)"))
beta_0_plot_m <- par_plot_mass(mass_fix_res, "beta_0", TeX(r"($beta_0$)"))
beta_f_plot_m <- par_plot_mass(mass_fix_res, "beta_f", TeX(r"($beta_f$)"))
beta_d_plot_m <- par_plot_mass(mass_fix_res, "beta_d", TeX(r"($beta_d$)"))
beta_1_plot_m <- par_plot_mass(mass_fix_res, "beta_1", TeX(r"($beta_1$)"))
beta_2_plot_m <- par_plot_mass(mass_fix_res, "beta_2", TeX(r"($beta_2$)"))
```


Arrange the parameter plots together.

```{r mass_inc_par, fig.height = 5, fig.width = 7, cache=TRUE}
ggarrange(sigma_w_plot_m, sigma_a_plot_m, alpha_plot_m, beta_0_plot_m,
          beta_f_plot_m, beta_d_plot_m, beta_1_plot_m, beta_2_plot_m,
          nrow=2, ncol=4, common.legend = TRUE,
          labels = LETTERS[1:8],
          font.label = list(size = 10),
          hjust = -1.2, vjust = 2,
          align ="hv") 
```


Both RMSE plots together:

```{r main_text_RMSE, fig.height = 4, fig.width = 7, cache=TRUE}
ggarrange(RMSE_plot, RMSE_plot_mass,
          nrow=1, ncol=2,
          labels = LETTERS[1:2],
          font.label = list(size = 10),
          hjust = -1.2, vjust = 8,
          align ="hv") 
```

# R info

```{r r_info}
sessionInfo()
```

## References

Auger-Méthé, M., C. Field, C. M. Albertsen, A. E. Derocher, M. A. Lewis, I. D. Jonsen, J. Mills Flemming.  2016. State-space models’ dirty little secrets: even simple linear Gaussian models can have estimation problems. Scientiic Reports 6: 26677. 10.1038/srep26677

Auger-Méthé, M., K. Newman, D. Cole, F. Empacher, R. Gryba, A. A. King, V. Leos-Barajas, J. Mills Flemming, A. Nielsen, G. Petris, and L. Thomas. 2021. A guide to state–space modeling of ecological time series. Ecological Monographs 91:e01470. 10.1002/ecm.1470 
````
