## Supplementary material for "Using wingbeat frequency to estimate mass gained by seabirds": wbf_mass_model.cpp

```
// Link to the TMB package
#include <TMB.hpp>

// Define main function
template<class Type>
Type objective_function<Type>::operator() ()
{
  // Specify the input data
  DATA_VECTOR(w);// Vector of wing beat frequency (WBF) values. Includes NAs when not in flight
  DATA_VECTOR(d); // Vector of 0/1 indicating when birds are diving/foraging
  DATA_VECTOR(f); // Vector of 0/1 indicating when birds are flying
  DATA_VECTOR(c1); // Vector of covariate 1
  DATA_VECTOR(c2); // Vector of covariate 2
  
  // bid is a matrix with 3 columns.
  // The first column (0) is the bird id value. There should be one row per bird.
  // The second column (1) is the index of the first observation for that individual
  // The third column (2) is the index of the last observation for that individual
  DATA_MATRIX(bid); 
  
  // For one-step-ahead residuals
  DATA_VECTOR_INDICATOR(keep, w);
  
  // Specify the parameters
  PARAMETER(logA); // mean population value for constant relating WBF and mass (>0) (log, alpha)
  PARAMETER(logSDP); // log of standard deviation for the process error (> 0) (sigma_w)
  PARAMETER(logSDO); // log of standard deviation for the observation error (>0) (sigma_m)
  PARAMETER(logB0);// base rate of mass loss (log)
  PARAMETER(logBf); // mass loss increase in flight (log)
  PARAMETER(bd); // intercept-like coefficient for mass gain while diving 
  PARAMETER(logSDA); // log of standard deviation for the random effect on a [constant relating mass to WBF] (>0)
  PARAMETER(b1); // coefficient for effect of covariate 1 on mass gain while diving
  PARAMETER(b2); // coefficient for effect of covariate 2 on mass gain while diving
  
  // Specify the random effect/states 
  PARAMETER_VECTOR(m); // mass
  PARAMETER_VECTOR(ai); // a for each individual

  // Vector for wpred
  vector<Type> wpred(w.size());

  // exp(par) is a trick to make sure that the estimates are > 0
  Type a = exp(logA);
  Type sdp = exp(logSDP);
  Type sdo = exp(logSDO);
  Type b0 = exp(logB0);
  Type bf = exp(logBf);
  Type sda = exp(logSDA);
  
  // Define the variable that will keep track of
  // the negative log-likelihood (nll)
  Type nll = 0.0;

  for(int b = 0; b < bid.rows(); ++b){
    
    // Calculate the contribution to the negative log-likelihood
    // of the process equation for t=1,...,T
    for(int i = (CppAD::Integer(bid(b,1))+1); i < (CppAD::Integer(bid(b,2))+1); ++i){
      nll -= dnorm(m(i), m(i-1) - b0 - (f(i-1) * bf) + (d(i-1) * exp(bd + (b1 * c1(i-1)) + (b2 * c2(i-1)))) , sdp, true);
      
      // Simulation block for process equation
      SIMULATE {
        m(i) = rnorm( m(i-1) - b0 - (f(i-1) * bf) + (d(i-1) * exp(bd + (b1 * c1(i-1)) + (b2 * c2(i-1)))), sdp);
      }
      
    }
    
    // Add random effect for constant influencing the relationship between mass and WBF for each bird
    nll -= dnorm(ai(b), a, sda, true); // ai is specific to each bird. It's assumed to be normally distributed around a
    SIMULATE {
      ai(b) = rnorm(a, sda);
    }
    
    // Calculate the contribution to the negative log-likelihood
    // of the observation equation for t=1,...,T
    for(int i = (CppAD::Integer(bid(b,1))+1); i < CppAD::Integer(bid(b,2)); ++i){
      if (w(i) == w(i)) { // ignore timesteps when fly is NA
        nll -= keep(i)*dnorm(w(i), ai(b)*sqrt(m(i)), sdo, true);
        wpred(i) = ai(b)*sqrt(m(i));
        
        // Simulation block for observation equation
        SIMULATE {
          w(i) = rnorm(ai(b)*sqrt(m(i)), sdo);
        }
        
      } else { //set the non-flying values of ypred to 0
        wpred(i) = w(i); 
      }
        
    }
  }
 
  // State the transformed parameters to report
  // Using ADREPORT will return the point values and the standard errors
  // Note that we only need to specify this for parameters
  // The other parameters, including the random effects (states),
  // will be returned automatically
  ADREPORT(sdo);
  ADREPORT(b0);
  ADREPORT(bf);
  ADREPORT(bd);
  ADREPORT(b1);
  ADREPORT(b2);
  ADREPORT(sdp);
  ADREPORT(a);
  ADREPORT(sdo);
  ADREPORT(sda);

  REPORT(wpred);
  REPORT(m);
  REPORT(ai);

  // Report simulated values
  SIMULATE{
    REPORT(m);
    REPORT(w);
    REPORT(ai);
  }
  
  // State that we want the negative log-likelihood to be returned
  // This is what the optimizer in R will minimize
  return nll;
}
```
